# Discovery and validation of novel human genomic safe harbor sites for gene and cell therapies

**DOI:** 10.1101/2021.03.04.433856

**Authors:** Erik Aznauryan, Alexander Yermanos, Elvira Kinzina, Edo Kapetanovic, Denitsa Milanova, George M. Church, Sai T. Reddy

## Abstract

Existing approaches for the integration and expression of genes of interest in a desired human cellular context are marred by the safety concerns related to either the random nature of viral-mediated integration or unpredictable pattern of gene expression in currently employed targeted genomic integration sites. Disadvantages of these methods lead to their limited use in clinical practice, thus encouraging future research in identifying novel human genomic sites that allow for predictable and safe expression of genes of interest. We conducted a bioinformatic search followed by experimental validation of novel genomic sites and identified two that demonstrated stable expression of integrated reporter and therapeutic genes without detrimental changes to cellular transcriptome. The cell-type agnostic criteria used in our bioinformatic search suggest wide-scale applicability of our sites for engineering of a diverse range of tissues for therapeutic as well as enhancement purposes, including modified T-cells for cancer therapy and engineered skin to ameliorate inherited diseases and aging. Additionally, the stable and robust levels of gene expression from identified sites allow for their use in industry-scale biomanufacturing of desired proteins in human cells.

## Introduction

Development of technologies for predictable, durable and safe expression of desired genetic constructs (i.e., transgenes) in human cells will contribute significantly to the improvement of gene and cell therapies (Bestor, 2000; Ellis, 2005), as well as for protein manufacturing (Lee et al., 2019). One prominent beneficiary of such technologies are genetically engineered T-cell therapies, which requires genomic integration of transgenes encoding novel immune receptors (Chen et al., 2020; Richardson et al., 2019); another example are gene therapies for highly proliferating tissues, such as inherited skin disorders, in which entire wild-type gene copies have to be integrated into epidermal stem cells (Droz-Georget Lathion et al., 2015; Hirsch et al., 2017). Advances in genome editing using targeted integration tools (Maeder and Gersbach, 2016) already allow precise genomic delivery and sustained expression of transgenes in certain cellular contexts, such as chimeric antigen receptors (CARs) integrated into the T cell receptor alpha chain locus in T-cells (Eyquem et al., 2017), and coagulation factors delivered to hepatocytes using recombinant adeno-associated viral (rAAV) vectors (Barzel et al., 2015). These applications, however, are limited to specific cell types and cause disruption to the endogenous genes, limiting the diversity of cellular engineering applications. Specific loci in the human genome that support stable and efficient transgene expression, without detrimentally altering cellular functions are known as Genomic Safe Harbor (GSH) sites. Thus, precise integration of functional genetic constructs into GSH sites greatly enhances genome engineering safety and efficacy for clinical and biotechnology applications.

Empirical studies have identified three sites that support long-term expression of transgenes: AAVS1, CCR5 and hRosa26 – all of which were established without any a-priori safety assessment of the genomic loci they reside in (Papapetrou and Schambach, 2016). The AAVS1 site, located in an intron of PPP1R12C gene region, has been observed to be a region for rare genomic integration events of the Adeno-associated virus’s payload (Oceguera-Yanez et al., 2016). Despite being successfully implemented for durable transgene expression in numerous cell types (Hong et al., 2017), the AAVS1 site location is in a gene-dense region, suggesting potential disruption of expression profiles of genes located in the vicinity of this loci (Sadelain et al., 2012). Additionally, studies indicated frequent transgene silencing and decrease in growth rate following transgene integration into AAVS1 (Ordovás et al., 2015; Shin et al., 2020), which represents a liability for clinical gene therapy. The second site lies within the CCR5 gene, which encodes a protein involved in chemotaxis and also serves as co-receptor for HIV cellular entry in T cells (Jiao et al., 2019). Serendipitously, researchers identified that the naturally-occurring CCR5-delta-32 mutation present in people of Scandinavian-origin results in an HIV-resistant phenotype (Silva and Stumpf, 2004). This finding suggested disposability of this gene and applicability of CCR5 locus for targeted genome engineering, especially for T cell therapies (Lombardo et al., 2011; Sather et al.). However, similar to AAVS1, the CCR5 locus is located in a gene-rich region, surrounded by tumor associated genes (Sadelain et al., 2012), thus severely limiting its safe use for therapeutic purposes. Additionally, CCR5 expression has been associated with promoting functional recovery following stroke (Joy et al., 2019), thus disrupting CCR5 may be undesirable in clinical practice. The third site, human Rosa26 (hRosa26) locus, was computationally predicted by searching the human genome for orthologous sequences of mouse Rosa26 (mRosa26) locus (Irion et al., 2007). The mRosa26 was originally identified in mouse embryonic stem cells by using random integration by lentiviral-mediated delivery of gene trapping constructs consisting of promotorless transgenes (β-galactosidase and neomycin phosphotransferase), resulting in sustainable expression of these transgenes throughout embryonic development (Friedrich and Soriano, 1991; Zambrowicz et al., 1997). Similar to the other two currently employed GSH sites, hRosa26 is located in an intron of a coding gene THUMPD3 (Irion et al., 2007), the function of which is still not fully characterized. This site is also surrounded by proto-oncogenes in its immediate vicinity (Sadelain et al., 2012), which may be upregulated following transgene insertion, thus potentially limiting the use of hRosa26 in clinical settings.

Attempts have been made to identify new human GSH sites that would satisfy various safety criteria, thus avoiding the disadvantages of existing sites. One approach developed by Sadelain and colleagues used lentiviral transfection of beta-globin and green fluorescence protein (GFP) genes into induced pluripotent stem cells (iPSCs), followed by the assessment of the integration sites in terms of their linear distance from various coding and regulatory elements in the genome, such as cancer genes, miRNAs and ultraconserved regions (Papapetrou et al., 2011). They discovered one lentiviral integration site that satisfied all of the proposed criteria, demonstrating sustainable expression upon erythroid differentiation of iPSCs. However, global transcriptome profile alterations of cells with transgenes integrated into this site were not assessed. A similar approach by Weiss and colleagues used lentiviral integration in Chinese hamster ovary (CHO) cells to identify sites supporting long-term protein expression for biotechnological applications (e.g., recombinant monoclonal antibody production) (Gaidukov et al., 2018). Although this study led to the evaluation of multiple sites for durable, high-level transgene expression in CHO cells, no extrapolation to human genomic sites was determined. Another study aimed at identifying novel GSHs through bioinformatic search of mCreI sites residing in loci that satisfy GSH criteria (Pellenz et al., 2019). Similarly, to previous work, several stably expressing sites were identified and proposed for synthetic biology applications in humans. However, local and global gene expression profiling following integration events in these sites have not been carried out.

All of the potential new GSH sites possess a shared limitation of being narrowed by lentiviral- or Cre-based integration mechanisms. Additionally, safety assessments of some of these newly identified sites, as well as previously established AAVS1, CCR5 and Rosa26, were carried out by evaluating the differential gene expression of genes located solely in the vicinity of these integration sites, without observing global transcriptomic changes following integration. A more comprehensive bioinformatic-guided and genome-wide search of GSH sites based on established criteria, followed by experimental assessment of transgene expression durability in various cell types and safety assessment using global transcriptome profiling would, thus, lead to the identification of a more reliable and clinically useful genomic region.

In this study, we used bioinformatic screening to rationally identify multiple sites that satisfy established as well as newly introduced GSH criteria. We then used CRISPR/Cas9 targeted genome editing to individually integrate a reporter gene into these sites to monitor long-term expression of the transgene in HEK293T and Jurkat cells. This experimental evaluation in cell lines was followed by testing of two promising candidate sites in primary human T-cells and human dermal fibroblasts using reporter and therapeutic transgenes, respectively. Finally, bulk and single-cell RNA-sequencing experiments were performed to analyze the transcriptomic effects of such integrations into these two newly established GSH sites.

## Results

### Bioinformatic search of novel GSH sites

To identify novel sites that could serve as potential GSHs, we first conducted a genome-wide bioinformatic search based on previously established and widely accepted (Sadelain et al., 2012) as well as newly introduced criteria that would satisfy safe and stable gene expression (Fig. 1A,B). We started by eliminating gene-encoding sequences and their flanking regions of 50 kb to thus avoid disruption of functional regions of gene expression. We then identified oncogenes and eliminated regions of 300 kb upstream and downstream to prevent insertional oncogenesis, a complication of gamma-retroviral and lentiviral integrations that may arise through unintended upregulation of an oncogene in the vicinity of the integration site (Hacein-Bey-Abina et al., 2008). We used oncogenes from both tier 1 (extensive evidence of association with cancer available) and tier 2 (strong indications of the association exist) to decrease the likelihood of oncogene activation upon integration. Additionally, genes can be substantially regulated by mircoRNAs, which cleave and decay mature transcripts as well as inhibit translation machinery, thus modulating protein abundance (Filipowicz et al., 2008). We, therefore, excluded miRNA-encoding regions and 300 kb long regions around them. Apart from promoters and microRNAs, gene expression may depend on the presence of enhancers that could be located kilobases away (Schoenfelder and Fraser, 2019; Vangala et al., 2020). We therefore excluded enhancers as well 20 kb regions around them, which provides an overall distance of up to 70 kb from gene-enhancer units, decreasing the chance of altering physiological gene expression. Additionally, we excluded regions surrounding long non-coding RNAs and tRNAs as well as 150 kb around them as they are involved in differentiation and development programs determining cell fate and are essential for normal protein translation, respectively (Guttman et al., 2009; Chen et al., 2016; Schimmel, 2018). Finally, we excluded centromeric and telomeric regions to prevent alterations in DNA replication, cellular division and normal aging (Villasante et al., 2007).

**Figure 1.**
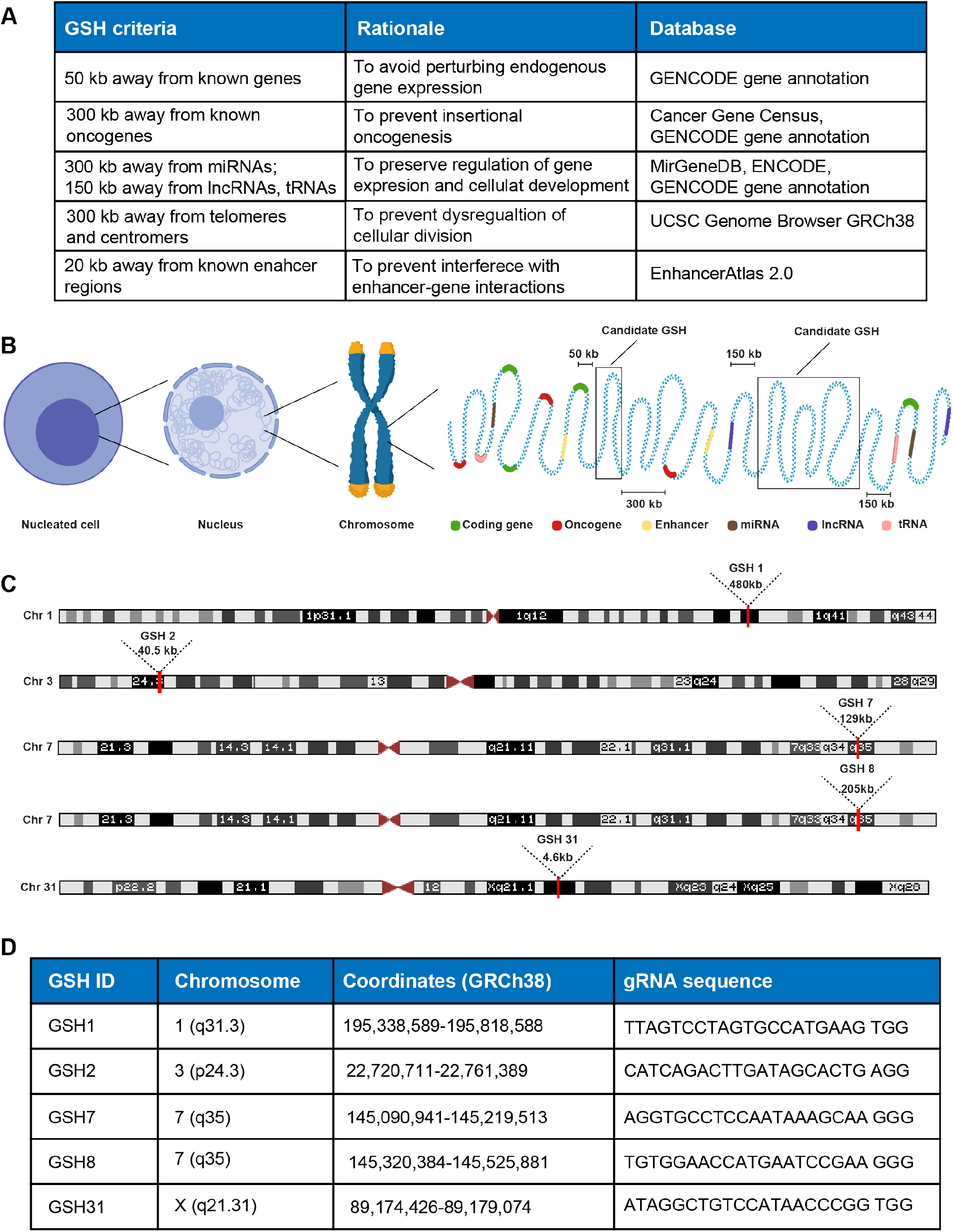
Bioinformatic identification of novel genomic safe harbor sites. A) Table shows GSH criteria, rationale and databases used to computationally predict GSH sites in the human genome. B) Schematic representation of candidate GSH sites, showing linear distances from different encoding and regulatory elements in the genome according to the established and newly introduced criteria. C) Chromosomal locations and lengths of five candidate GSH sites, which were subsequently experimentally tested. D) Chromosomal coordinates of five candidate GSH sites and the gRNA sequences used for subsequent CRISPR/Cas9 genome editing. See also Supplementary table 1 for the list of all computationally predicted sites.

Based on our bioinformatic screening, we identified close to two thousand sites that satisfied all of our criteria (Sup. table 1). We chose five sites that varied significantly in size (GSH1, 2, 7, 8, GSH31), designed guide RNAs (gRNA) targeting these sites and possessing high on- and off-target scores (high on-target and low off-target activities), and characterized the durability and safety of transgene expression at these sites experimentally (Fig. 1C,D).

### Experimental validation of bioinformatically identified GSH sites by targeted transgene integration in human cell lines

In order to experimentally assess transgene expression from the five predicted novel GSH sites, we performed targeted integration of a gene construct encoding a red fluorescence reporter protein (mRuby) into two common human cell lines – HEK293T and Jurkat cells. HEK293 are commonly used for medium- to large-scale production of recombinant proteins (Chin et al., 2019), thus identifying GSH in HEK293 may be relevant for protein manufacturing. The Jurkat cell line was derived from T-cells of a pediatric patient with acute lymphoblastic leukemia (Abraham and Weiss, 2004) and has been used extensively used for assessing the functionality of engineered immune receptors, thus discovery of GSH in this cell line supports applications in T cell therapies (Roybal et al., 2016; Vazquez-Lombardi et al., 2020). For integration of mRuby, we employed a CRISPR/Cas9-based genome editing strategy that uses the Precise Integration into Target Chromosome (PITCh) method (Nakade et al., 2014; Sakuma et al., 2016), assisted by microhomology-mediated end-joining (MMEJ) (Sfeir and Symington, 2015). This approach utilizes a reporter-bearing plasmid possessing short microhomology sequences flanked by gRNA binding sites. Once inside the cell the reporter gene together with microhomologies directed against the candidate GSH site are liberated from the plasmid by Cas9-generated double-stranded breaks (DSB) at gRNA binding sites on the PITCh donor plasmid. A different gRNA-Cas9 pair generates DSBs at the candidate GSH locus, and the freed reporter gene with flanking micro-homologies is integrated by exploiting the MMEJ repair pathway (Fig. 2A,B). This PITCh MMEJ approach allowed us to rapidly generate donor plasmids targeted against different predicted safe harbor sites, in contrast to the more elaborate process of cloning long homology arms (i.e., >300 bp) required for homology-directed repair (HDR). The error-prone mechanism of MMEJ-mediated integration did not represent a substantial concern since the targeted sites are distanced from any identified coding or regulatory element and thus mutations arising following integration are unlikely to cause any detrimental changes.

**Figure 2.**
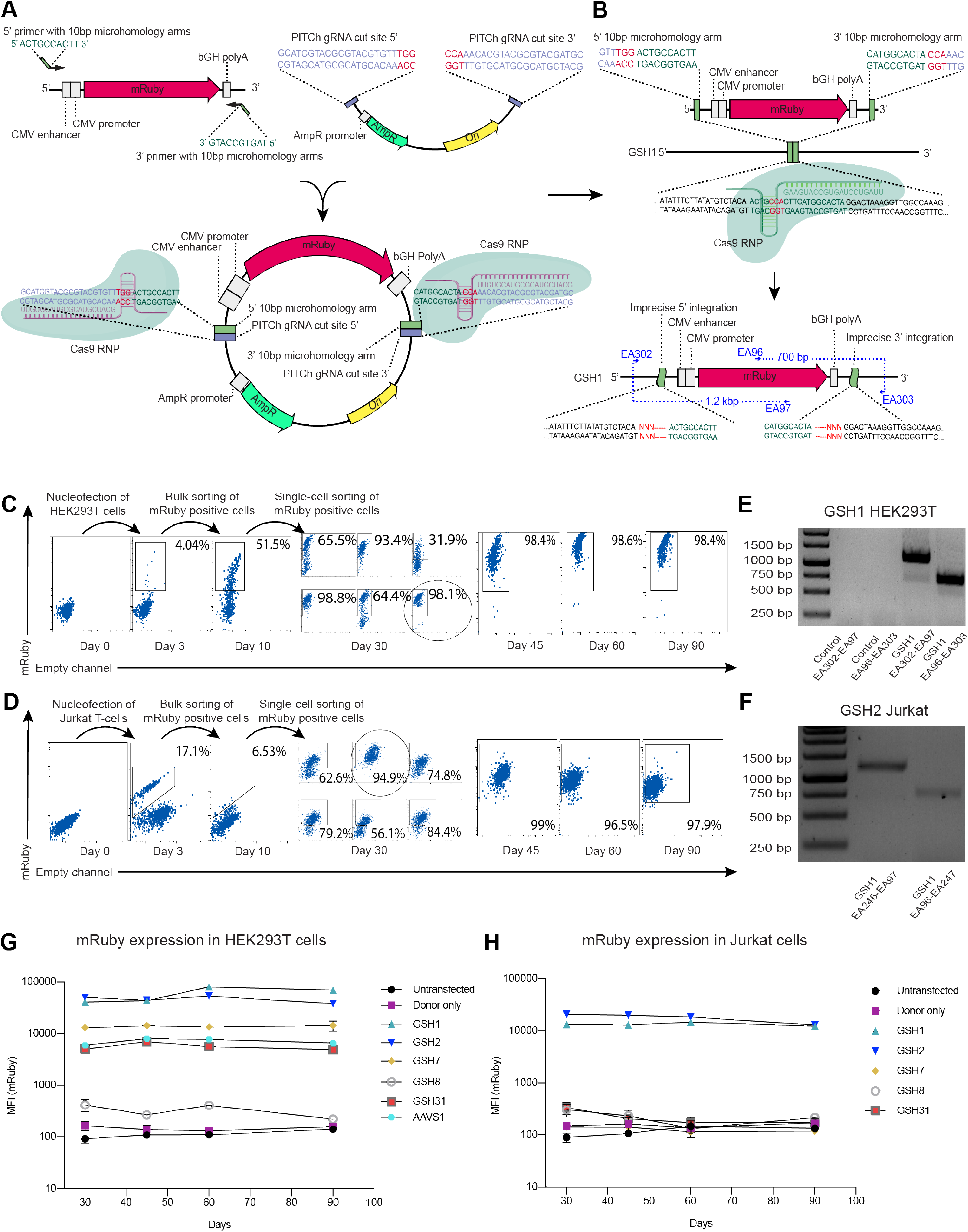
Experimental validation of candidate GSH sites by targeted genome editing in HEK293T and Jurkat cells. A) PITCh plasmid is generated by cloning an mRuby-bearing insert with micro-homologies against specific GSH into a backbone possessing PITCh gRNA target sites, needed for the liberation of the insert inside the engineered cell by Cas9. B) Once inside the cell, the mRuby insert is integrated into a desired site by the MMEJpathway following a Cas9-induced double-stranded break of the targeted site. C, D) Flow cytometry demonstrating the isolation of clonal populations expressing the mRuby transgene from GSH1 locus in HEK293T cells and GSH2 locus in Jurkat cells using pooled and single-cell flow cytometry mediated sortings. The highest expressing GSH1-HEK293T clone and GSH2-Jurkat clone was expanded in cell culture and flow cytometry measurements at day 45, 60 and 90 demonstrated stable levels of transgene expression. E, F) Genotyping of the GSH1 site in HEK293T cells and GSH2 site in Jurkat cells using primers spanning the junction between integration site and the trangene show mRuby integration into the predicted locus. G) mRuby transgene integration into each of the tested GSH sites in HEK293T show stable expression from GSH1, GSH2, GSH7 and GSH 31. Data are represented as mean ± SEM, N=2. H) mRuby transgene integration into each of the tested GSH sites in Jurkat show stable expression from GSH1 and GSH2. Data are represented as mean ± SEM, N=2.

Using the PITCh approach, we transfected mRuby transgene into the five candidate GSH sites using the best predicted gRNA sequence for each site (see Methods). We then conducted a pooled selection of mRuby-expressing HEK293T and Jurkat cells by fluorescence-activated cell sorting (FACS), followed by expansion for one week and single-cell sorting to produce monoclonal populations of mRuby-expressing cells. In order to determine sites that support long-term stable transgene expression, we monitored clones with homogenous and high mRuby expression levels by performing flow cytometry at day 30, 45, 60 and 90 after integration.

Out of five candidate GSH sites, four sites in HEK293T cells – GSH1, 2, 7 and 31 (Fig. 2C,G) – and two sites in Jurkat cells – GSH1 and 2 (Fig. 2D,H) – demonstrated stable mRuby expression levels 90 days after integration. Interestingly, two sites in HEK293T cells – GSH1, GSH2 – allowed for over an order of magnitude higher transgene expression levels as compared to the commonly used AAVS1 site throughout the 90-day duration of cell culture (Fig. 2G). Transgene integration into these sites was confirmed by genotyping using primer pairs amplifying the junction between tested GSH and the transgene (Fig. 2E,F).

### Transcriptome profiling of cell lines following targeted integration in GSH sites

In order to assess whether targeted integration into the candidate GSH sites resulted in aberration of the global transcriptome profiles, we performed a bulk RNA-sequencing and analysis. Following ninety days in culture the clone showing the highest GSH2-integrated mRuby levels was compared with untreated cells from the same culture for both HEK293T and Jurkat cells (Fig. 3A). Paired-end sequencing on Ilumina NextSeq500 with an average read length of 100 base-pairs and 30 million reads per sample was employed on two biological replicates of untreated and GSH2-mRuby cultures of HEK293T and Jurkat cells. We first performed a principal component analysis and visualized each sample in two-dimensions using the first two principal components. This immediately revealed transcriptional similarity within the integrated and wild-type samples of the same biological replicate for both cell lines (Fig. 3B). While biological variation was observed between the HEK293T samples, the Jurkat samples, both treated and untreated, maintained conserved transcriptional profiles. Performing differential gene expression analysis revealed minor differences between integrated and unintegrated samples for both cell lines relative to the differences between the two cell types (Fig. 3C). It was additionally promising that the most differentially expressed genes were not shared between Jurkat and HEK293T cell lines, further suggesting integration in GSH2 does not systematically alter gene expression. Interestingly, differentially expressed genes were scattered across different chromosomes, as opposed to being concentrated within the integrated chromosome where more local contacts exist, again pointing at biological variation (Figure 3D). Furthermore, performing gene ontology analysis revealed no significant enrichment of cancer associated genes or pathways in both HEK and Jurkat cells (Fig. S1, S2), again supporting the potential safety of the GSH2 site. We lastly quantified the differences in gene expression for both cell lines either across biological replicates without GSH2 integration versus within a biological replicate with or without GSH2 integration (Fig. 3E). Mirroring our principal component analysis (Fig. 3B), this analysis again supports that the differences in gene expression we observe arise from biological variation between clones and not due to integration at GSH2.

**Figure 3.**
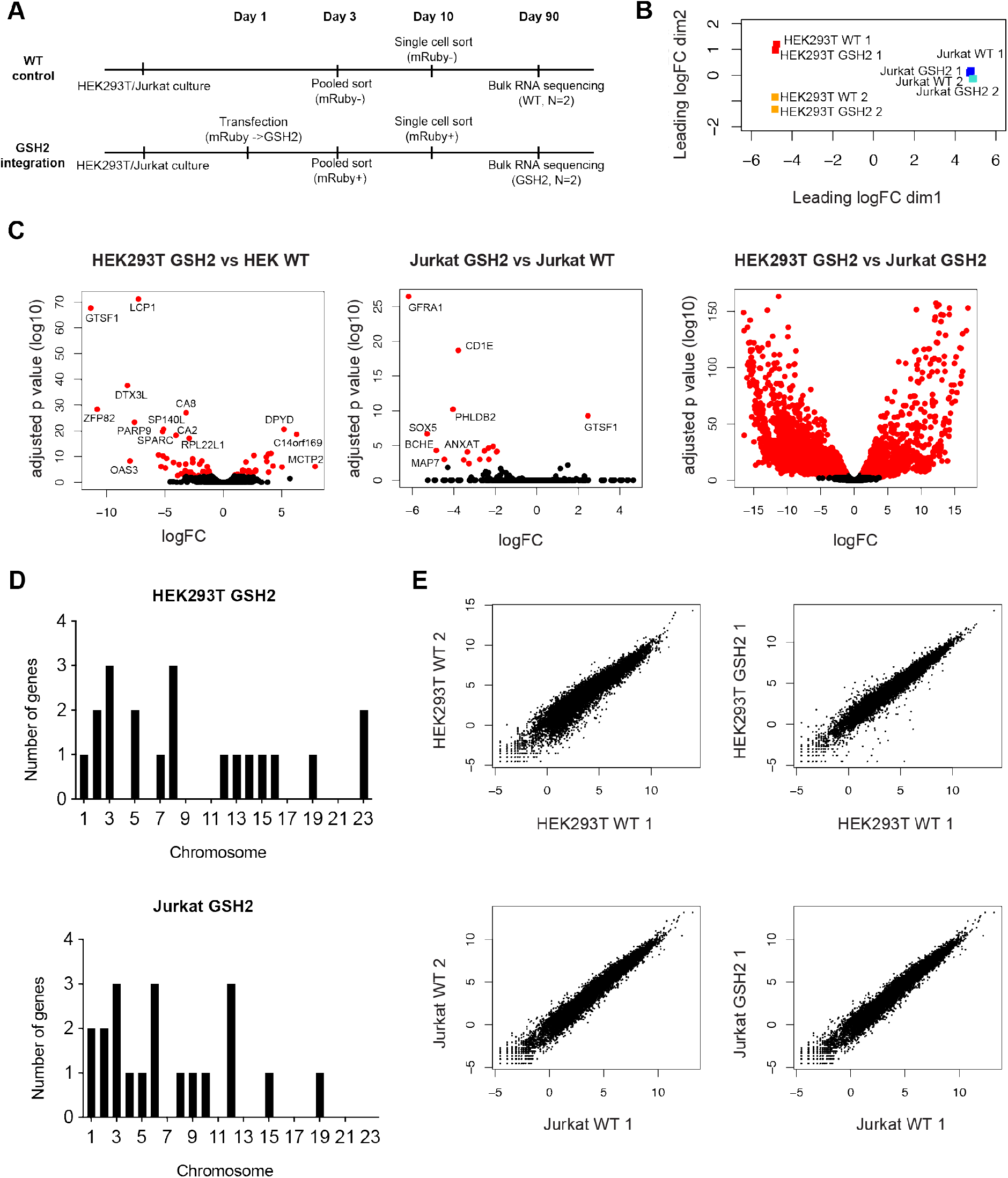
RNA sequencing and transcriptome analysis of HEK293T and Jurkat cells following mRuby integration into GSH2. A) Pipeline of bulk RNA-seq experiment on GSH2 integrated and non-integrated HEK293T and Jurkat cells. B) PCA of two biological replicates of HEK293T and Jurkat cells with and without mRuby integration into GSH2. C) Differential expression of genes following GSH2 integration in HEK293T and Jurkat and comparison of HEK293T and Jurkat non-integrated cells. D) Chromosomal distribution of differentially expressed genes in HEK293T and Jurkat cells. Genes with an adjusted p-value of less than 0.05 were considered differentially expressed. E) Correlation of gene expression either between biological replicates without GSH2 integration or within a biological replicate with or without integration in GSH2. See also S1 and S2 for the functional classification of differentially expressed genes in HEK and Jurkat, respectively.

### Targeted integration in novel GSH sites in primary human T-cells and primary human dermal fibroblasts

We next sought to characterize targeted integration into GSH1 and GSH2 sites in primary human cells. One of the potential applications of targeted integration into novel GSH sites is for the ex-vivo engineering of human T-cells, which are being extensively explored for adoptive cell therapies in cancer and autoimmune disease. Thus, we first tested GSH1 and GSH2 in primary human T-cells isolated from peripheral blood of a healthy donor. This time we targeted these sites by employing an HDR-based integration approach using a linear double-stranded DNA donor template, which contained the mRuby transgene driven by a CMV promoter and with 300bp homology arms (Fig. 4A). Phosphorothioate bonds and biotin groups were also added to 5’ and 3’ ends of the HDR template to increase its stability and prevent concatemerization, respectively (Gutierrez-Triana et al., 2018). Nucleofection of Cas9-gRNA ribonucleoprotein (RNP) complexes and HDR templates into primary T-cells resulted in mRuby-positive expression in 1.3% of cells for GSH1 and 1.24% of cells for GSH2. These mRuby-expressing cells were isolated by FACS on day four, cultured for another seven days; a second round of sorting was performed on the mRuby-positive populations. Following these two rounds of pooled sorting, a highly enriched population of T cells stably expressing the mRuby transgene was isolated and cultured for the duration of T cell ex-vivo culture (up to day 20), with mRuby expression from GSH1 and GSH2 in 94.7% and 91.8% of cells, respectively (Fig. 4B). Correct integration into GSH1 and GSH2 was confirmed by genotyping and Sanger-sequencing using primers amplifying the junction between GSH1/GSH2 loci and the mRuby donor (Fig. 4C).

**Figure 4.**
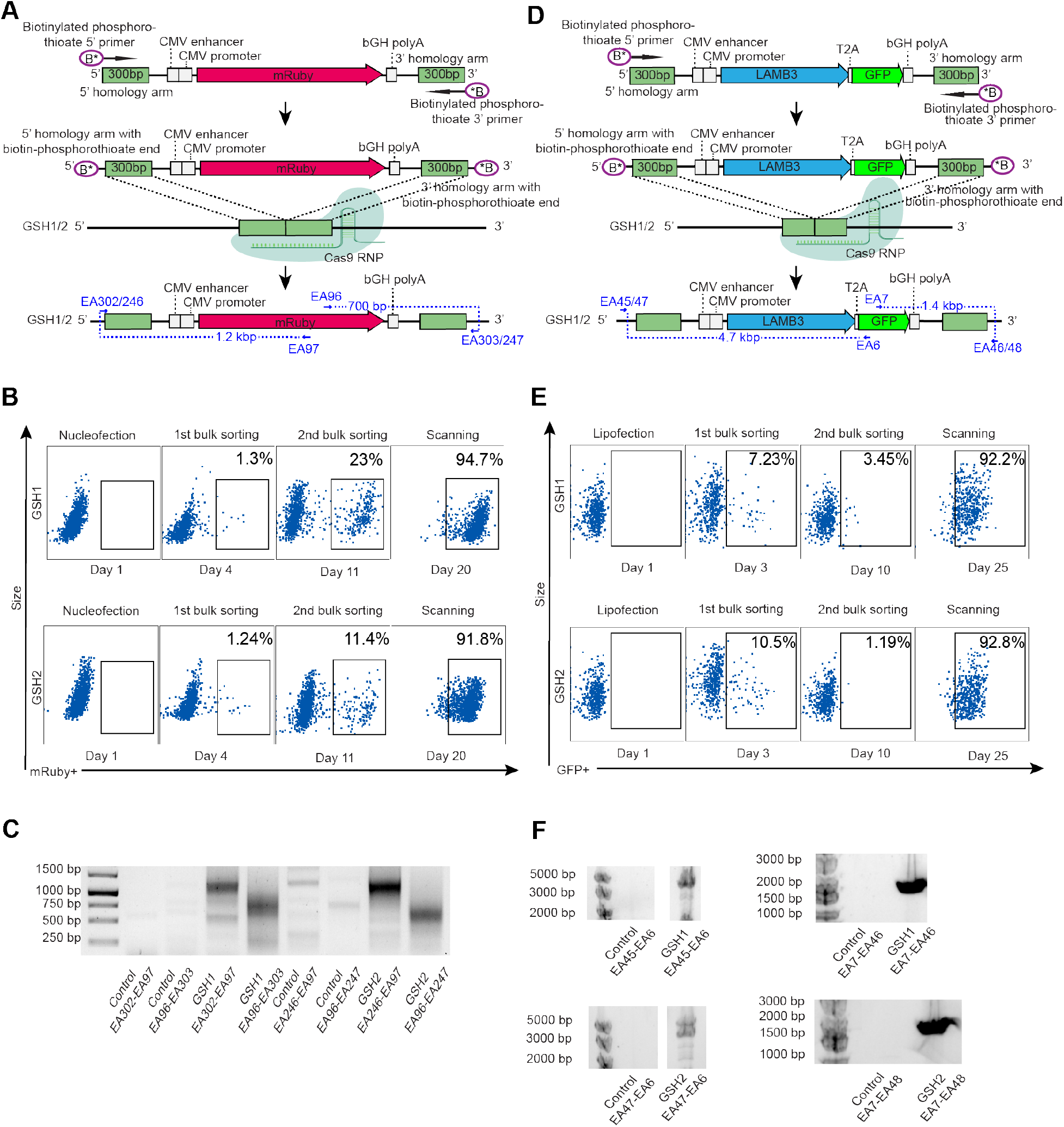
Targeted transgene integration into GSH1 and GSH2 in primary human cells. A) Targeted integration of mRuby into GSH1 and GSH2 in primary human T cells by Cas9 HDR. B) Flow cytometry plots demonstrating mRuby expression in both GSH1 and GSH2 in primary human T cells following two rounds of pooled sorting. C) PCR-based genotyping of GSH1 and GSH2 sites by using primers spanning the junction of targeted site and the inserted transgene indicate correct integration of mRuby in primary human T cells. D) Targeted integration of LAMB3-T2A-GFP into GSH1 and GSH2 in primary human dermal fibroblasts by Cas9 HDR. E) Flow cytometry plots demonstrating GFP expression in both GSH1 and GSH2 in primary human dermal fibroblasts following two rounds of pooled sorting. F) PCR-based genotyping of GSH1 and GSH2 sites by using primers spanning the junction of targeted site and the inserted transgene indicate correct integration of LAMB3-T2A-GFP in primary human dermal fibroblasts. See also Supplementary table 2 for precise sequences of donor constructs.

Another possible ex-vivo application of identified GSH sites includes engineering dermal fibroblasts and keratinocytes for autologous skin grafting in people with burns or inherited skin disorders. A group of genetic skin disorders named junctional epidermolysis bullosa (JEB) is associated primarily with mutations in a family of multi-subunit laminin proteins, which are involved in anchoring the epidermis layer of the skin to derma (Bardhan et al., 2020). Certain variants of JEB are specifically related to mutations in a beta subunit of laminin-5 protein, encoded by the *LAMB3* gene (Robbins et al., 2001). Using a similar dsDNA HDR donor with 300bp homology arms possessing phosphorothioate bond and biotin, we used Cas9 HDR to integrate the *LAMB3* gene tagged with GFP (total insert size 5,409 bp) into GSH1 and GSH2 sites in primary human dermal fibroblasts isolated from neonatal skin (Fig. 4D). After lipofection of fibroblasts with Cas9 and HDR templates, expression of GFP, which is indicative of LAMB3 expression, was observed in 7.23% (GSH1) and 10.5% (GSH2) of cells. These cells were sorted at day three, cultured for seven days and the GFP-positive population – 3.45% for GSH1 and 1.19% for GSH2 – was sorted again. Similar to T-cells, two rounds of pooled sorting led to over 92% enrichment of GFP-positive cells, with the expression of *LAMB3*-GFP transgene maintained for the duration of cell culture (over 25 days) (Fig. 4E). Genotyping and Sanger-sequencing confirmed successful integration into both loci by using primers amplifying the junction between GSH1/GSH2 and the LAMB3-GFP donor (Fig. 4F).

### Single-cell RNA sequencing and analysis of primary human T cells following transgene integration into a novel GSH site

Lastly, we assessed transcriptome-wide effects on a single-cell level following transgene integration into GSH1 in primary T-cells. We performed single-cell RNA sequencing using the 10X Genomics protocol, which consists of encapsulating cells in gel beads bearing reverse transcription (RT) reaction mix with unique cell primers. Following the RT reaction, the cDNA is pooled, and the library is amplified for subsequent next-generation sequencing.

This single-cell sequencing workflow was applied to human T cells expressing mRuby in GSH1 after 25 days in culture, wildtype (non-transfected) cells were used as a control. We also compared these cells with wild-type controls from a different donor to again compare whether GSH integration resulted in more variability in gene expression relative to a biological replicate (Fig. 5A). Performing differential gene expression analysis across the three samples revealed fewer up- or downregulated genes following GSH1 integration relative to the untreated, second patient sample (Fig. 5B). We performed uniform manifold approximation projection (UMAP) paired with an unbiased clustering based on global gene expression, which resulted in 13 distinct clusters (Fig. 5C). Many genes defining these clusters corresponded to typical T cell markers such as IL7R, ICOS, CD28, CCL5, CD74, and NKG7 (Fig. 5D). We subsequently quantified the proportion of cells per cluster for each sample, again demonstrating congruent gene expression signatures from cells arising from a single patient, regardless of whether integration in GSH1 occurred or not (Fig. 5E). Furthermore, similar to bulk RNA-sequencing results on cell lines, none of the most differentially expressed genes that were upregulated in cells with GSH1 transgene integration were associated with any cancer-related pathways (Fig. 5F). Interestingly, the expression of the Jun gene encoding the oncogenic c-Jun transcription factor is decreased in cells bearing transgene integration into GSH1. Taken together, both our single-cell and bulk RNA-sequencing data suggest that the computationally determined and experimentally validated GSHs have minimal influences on global gene expression.

**Figure 5.**
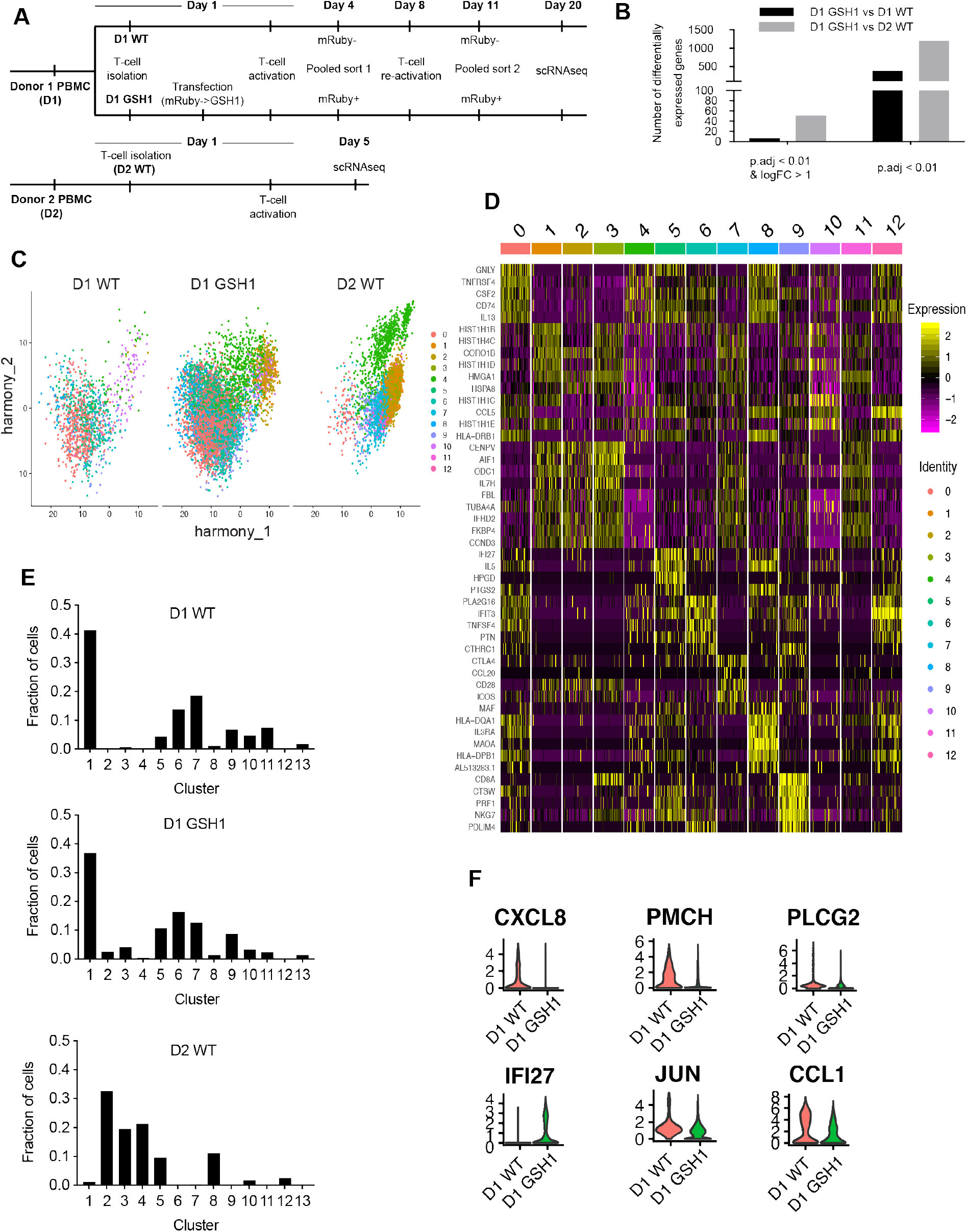
Single-cell RNA-seq of primary human T-cells following targeted transgene integration into GSH1 site. A) Pipeline of the RNA-seq experiment following Cas9 HDR targeted integration of mRuby into GSH1 (GSH1-mRuby cells) and T-cell activation. B) Number of differentially expressed genes GSH1-mRuby T-cells and WT T-cells (non-integrated) from donor 1 and GSH1-mRuby T-cells from donor 1 and WT T-cells from donor 2. C) UMAP analysis comparing transcriptional clusters of GSH1-mRuby and WT T-cells from donor 1 and WT T-cells from donor 2. Each point represents a unique cell barcode, and each color corresponds to cluster identity. D) Expression of genes determining the seven largest clusters. Intensity corresponds to normalized gene expression. E) Distribution of GSH1-mRuby-and WT T-cells from donor 1 and WT T-cells from donor 2 across different clusters. F) Normalized expression for selected differentially expressed genes between GSH1mRuby and WT T-cells from donor 1.

## Discussion

In this study we used bioinformatic screening to identify novel GSH sites and performed phenotypic validation by targeted transgene integration in human cell lines and primary cells, resulting in durable and stable transgene expression. The potential safety of GSH sites was confirmed by observing minimal changes in transcriptomic profiles following transgene integration. None of the upregulated transcripts following transgene integration were associated with any of the known cancer pathways. These findings make the newly identified sites potentially preferable to currently used AAVS1, CCR5 and hRosa26, which have the drawbacks of being located within functional genes, in gene-dense regions and surrounded by oncogenes (Sadelain et al., 2012). Although previous studies have also resulted in the discovery of sites capable of long-term expression of transgenes, they were limited by the integration mechanism researchers employed and changes to the entire transcriptome following integration events were not evaluated, as they were focused on differential expression of a handful of genes in the vicinity of the discovered site (Papapetrou et al., 2011). Finally, generalizability of the criteria used to establish our new GSH sites suggests their possible applicability to different cell types, expanding the genome engineering toolkit for diverse cell therapy and synthetic biology applications (Nielsen and Voigt, 2014).

The most immediate use of identified GSH sites may involve safe and predictable engineering of human T-cells for adoptive cell therapy applications (Schwarz and Leonard, 2016). Copious endeavors to design, modify and augment functions of T-cells ex-vivo have been successfully initiated in research labs (Baeuerle et al., 2019; Eyquem et al., 2017). However, most strategies have relied on viral-mediated delivery, which results in random transgene integration and is thus associated with the risk of insertional oncogenesis, potentially leading to cancerous transformations of engineered cells, and unpredictability of transgene expression levels associated with the nature of the integration locus and frequent silencing of the integrated construct. Performing targeted integration into GSH sites would enable long-term transgene expression in a safe manner and would support advanced efforts in engineered T cell therapies such as armored CAR-T cells, capable of overcoming hostile tumor microenviroments (Yeku et. al., 2017) as well as T cells bearing synthetic receptors that introduce logic gates into cell’s behavior, allowing for safer and more effective treatments (Roybal and Lim, 2017). Additionally, given the demonstrated efficiency in dermal fibroblasts, we envision a rapid application of the discovered sites to skin engineering, particularly in the context of treatment of the inherited skin disorders, wound healing as well as skin rejuvenation.

Another exciting aspect of the identified GSH sites is the level of transgene expression observed, especially in HEK293T cells, which are known to be suitable for large-scale production of therapeutic proteins. We observed high levels of reporter gene expression from GSH1 and GSH2 in HEK293T that were sustained for over three months and exceeded expression levels from the AAVS1 site. This high expression level can theoretically be enhanced further by multiple biallelic integration events into identified loci and thus be exploited for durable large-scale production of commercially valuable proteins.

In summary, two novel human genomic safe harbor sites identified and validated in this study may serve as a robust and safe platform for a variety of clinically and industrially relevant cell engineering approaches, culminating in safer and more reliable gene and cell therapies.

## Methods

### Computational search for GSH sites

Previously established criteria (Sadelain et al., 2012) as well as newly introduced ones were used to predict genomic locations of novel GSHs. Specifically, coordinates of all known genes were extracted from GENCODE gene annotation (Release 24). A set of tier 1 and tier 2 oncogenes was obtained from Cancer Gene Census. The miRNA coordinates were obtained from MirGeneDB (Fromm et al., 2020). Enhancer regions were obtained from the EnhancerAtlas 2.0 database (Gao and Qian, 2019), coordinates were transposed into GRCh38/hg38 genome and union of enhancer sites was used. Genomic locations of sequences of tRNA and lncRNA were extracted from GENCODE gene annotation (Release 24). UCSC genome browser GRCh38/hg38 was used to get coordinates of telomeres and centromeres as well as unannotated regions. BEDTools (Quinlan and Hall, 2010) were used to determine flanking regions of each element of the criteria as well as to obtain union or difference between sets of coordinates. The custom source code developed for the computational identification of novel human genomic safe harbor sites is available at https://github.com/elvirakinzina/GSH.

### Plasmids, guide RNA design and HDR donor generation

PITCh plasmids were generated through standard cloning methods. CMV-mRuby-bGH insert was amplified from pcDNA3-mRuby2 plasmid (Addgene, Plasmid #40260) with primers containing mircohomology sequences against specific GSHs and AAVS1 site with 10bp of overlapping ends for the pcDNA3 backbone. The pcDNA3 backbone was amplified with primers containing sequences of PITCh gRNA cut site (GCATCGTACGCGTACGTGTTTGG) on both 5’ and 3’ ends of the backbone. The insert and the backbone were assembled using Gibson Assembly Master Mix (New England Biolabs, #E2611L).

Guide RNA sequences for five tested GSH sites were predicted using Geneious gRNA design tool. Briefly, coordinates of the predicted GSH sites were pasted into UCSC Genome Browser GRCh38/hg38 and DNA sequences were extracted and transferred into Geneious. An internal gRNA design tool was used to identify gRNA sequences located in the predicted GSHs against the entire human genome. The evaluation of the efficacy of double-stranded break generation (on-target activity) was based on Doench et al., 2016, while the specificity of the gRNA-induced break (off-target activity) was assessed based on Hsu et al., 2013. Guide RNAs with high on-target and off-target scores were used to target predicted GSHs.

Plasmids encoding CMV-mRuby-bGH flanked by GSH1/GSH2 300bp homology arms were ordered from Twist Biosciences in pENTR vector. HDR donors were amplified from these plasmids using biotinylated primers with phosphorothioate bonds between the first 5 nucleotides on both 5’ and 3’ ends. Plasmid encoding CMV-LAMB3-T2A-GFP-bGH was generated by overlap extension PCR of LAMB3 cDNA, purchased from Genscript (NM_000228.3) and GFP-bGH sequence from Addgene (Plasmid #11154). T2A sequence was added to 5’primer of GFP-bGH. Produced insert was cloned into the abovementioned pENTR vector from Twist Biosciences bearing GSH1 and GSH2 300bp homology arms as well as CMV promoter sequence using Gibson Assembly Master Mix (NEB, #E2611L). HDR donors were amplified from these plasmids using biotinylated primers with phosphorothioate bonds between the first 5 nucleotides on both 5’ and 3’ ends. HDR donors were then purified from PCR mix using SPRI beads (Beckman Coulter, #B23318) at 0.4X beads to PCR mix ratio.

### HEK293T and Jurkat cell culture, transfection and sorting

HEK293T cells were obtained from the American Type Culture Collection (ATCC) (#CRL-3216); the Jurkat leukemia E6-1 T cell line was obtained from ATCC (#TIB152). HEK cells were cultured in Dulbecco’s Modified Eagle’s Medium (DMEM) (ATCC 30-2002) supplemented with 2mM L-glutamine (ATCC 30-2214). Jurkat cells were cultured in ATCC-modified RPMI-1640 (Thermo Fisher, #A1049101). All media were supplemented with 10% FBS, 50 U ml-1penicillin and 50 μg ml^−1^ streptomycin. Detachment of HEK cells for passaging was performed using the TrypLE reagent (Thermo Fisher, #12605010). All cell lines were cultured at 37°C, 5% CO2 in a humidified atmosphere.

Prior to transfection of HEK293T and Jurkat gRNA molecules were assembled by mixing 4 μl of custom Alt-R crRNA (200 μM, IDT) with 4 μL of Alt-R tracrRNA (200 μM, IDT, #1072534), incubating the mix at 95°C for 5 min and cooling it to room temperature. 2 μL of assembled gRNA molecules were mixed with 2 μL of recombinant SpCas9 (61 μM, IDT, #1081059) and incubated for > 10 min at room temperature to generate Cas9 RNP complexes.

For transfection of HEK cells 100 μL format SF Cell line kit (Lonza, V4XC-2012) and electroporation program CM-130 was used on the 4D-Nucleofector. 1×10^6^ HEK cells were transfected with 2 μg of PITCh donor, 2 μl of Cas9 RNP complex against specific GSH and 2 μl of Cas9 RNP complex against PITCh plasmid to liberate MMEJ insert.

For transfection of Jurkat cells 100 μL format SE Cell line kit (Lonza, V4XC-1012) and electroporation program CL-120 was used on the 4D-Nucleofector. 1×10^6^ Jurkat cells were transfected with 2 μg of PITCh donor, 2 μl of Cas9 RNP complex against specific GSH and 2 μl of Cas9 RNP complex against PITCh plasmid to liberate MMEJ insert.

Transfected HEK and Jurkat cells were bulk sorted on day 3 and single-cell sorted on day 10 following transfection using Sony SH800S sorter. Best expressing clone was selected on day 30, split into two wells and cultured for another 2 months. mRuby expression of the best expressing clone was analyzed on BD LSRFortessa Flow Cytometer on day 45, 60 and 90 following transfection.

### Human T-cells culture, transfection and sorting

Human peripheral blood mononuclear cells were purchased from Stemcell Technologies (#70025) and T cells isolated using the EasySep Human T Cell Isolation kit (Stemcell Technologies, #17951). Primary human T cells were cultured for up to 20 days in ATCC-modified RPMI (Thermo Fisher, #A1049101) supplemented with 10% FBS, 10 mM non-essential amino acids, 50 μM 2-mercaptoethanol, 50 U ml-1penicillin, 50 μg ml^-6^ streptomycin and freshly added 20 ng ml^−1^ recombinant human IL-2, (Peprotech, #200-02). T cells were cultured at 37°C, 5% CO2 in a humidified atmosphere. On day 1 of culture, transfection of primary T cells with Cas9 RNP complexes and GSH1/GSH2-mRuby HDR templates was performed using the 4D-Nucleofector and a 20 uL format P3 Primary Cell kit (Lonza, V4XP-3032). Briefly, gRNA molecules were assembled by mixing 4 μl of custom Alt-R crRNA (200 μM, IDT) with 4 μL of Alt-R tracrRNA (200 μM, IDT, #1072534), incubating the mix at 95°C for 5 min and cooling it to room temperature. 2 μL of assembled gRNA molecules were mixed with 2 μL of recombinant SpCas9 (61 μM, IDT, #1081059) and incubated for > 10 min at room temperature to generate Cas9 RNP complexes. 1×10^6^ primary T cells were transfected with 1 μg of HDR template, 1 μl of GHS1/GSH2 Cas9 RNP complex using the EO115 electroporation program. T cells were activated with Dynabeads™ Human T-Activator CD3/CD28 (Thermo Fischer, #11161D) 3-4 hours following transfection. mRuby-positive T-cells were bulk sorted on day 4 using Sony SH800S sorter, re-activated with the new beads on day 8, sorted again on day 11 and analyzed on BD LSRFortessa Flow Cytometer on day 20.

### Human dermal fibroblasts culture, transfection and sorting

Neonatal human dermal fibroblasts were purchased from Coriell Institute (Catalog ID GM03377). Primary fibroblasts were cultured for up to 25 days in Prime Fibroblast media (CELLNTEC, CnT-PR-F). Cells were passaged at 70% confluency using Accutase (CELLNTEC, CnT-Accutase-100). Detached cells were centrifuged for 5 min, 200 x g at room temperature and seeded at seeded at 2,000 cells per cm^2^. Fibroblasts were cultured at 37°C, 5% CO2 in a humidified atmosphere. Fibroblasts were transfected using Lipofectamine™ CRISPRMAX™ Cas9 Transfection Reagent (ThermoFisher Scientific, CMAX00001). Briefly, cells were transfected at 50% confluency with 1:1 ratio of custom sgRNA (40 pmoles, Synthego) and SpCas9 (40pmoles, Synthego) and 2.5 μg of GSH1/GSH2 LAMB3-T2A-GFP HDR template. GFP-positive fibroblasts were bulk sorted on day 3 and 10 using Sony SH800S sorter and analyzed on BD LSRFortessa Flow Cytometer on day 25.

### Genotypic analysis of GSH integration

Genomic DNA was extracted from 1×10^6^ cells using PureLink Genomic DNA extraction kit (ThermoFischer Scientific, #K1820-01). 5 μL of genomic DNA extract were then used as templates for 25 μL PCR reactions using a primer with one primer residing outside of the homology arm of the integrated sequence and the other primer inside the integrated sequence. Obtained bands were gel extracted using Zymoclean Gel DNA Recovery Kit (Zymo Research, #D4001), 4ul of eluted DNA was cloned into a TOPO-vector using Zero-blunt TOPO PCR Cloning Kit (ThermoFischer Scientific, #450245), incubated for 1 hour, transformed into NEB 5-alpha Competent E. coli cells (New England Biolabs, C2987H) and plated on agar plates containing kanamycin at 50 μg/ml. Produced clones were picked and inoculated for overnight culture in 5ml of liquid broth supplemented with kanamycin at 50 μg/ml. Liquid cultures were mini-prepped the following morning using ZR Plasmid Miniprep - Classic kit (Zymo Research, #D4015) and Sanger sequenced by Microsynth using M13-forward and M13-reverse standard primers.

### Bulk RNA-sequencing of HEK293T and Jurkat cells GSH2 and WT

Following single-cell sort, the best expressing clone (GSH2) and wild-type (WT) of HEK293T and Jurkat cells were split into 2 wells (1 and 2) and cultured for 80 days, after which total RNA was extracted using PureLink RNA Mini Kit (ThermoFischer Scientific, #12183018A). Extracted total RNA was depleted of rRNA using RiboCop rRNA Depletion Kit (Lexogen, #144), first and second strands of cDNA were generated with SuperScript Double-Stranded cDNA Synthesis Kit (ThermoFischer Scientific, #11917010) using random hexamers and flow cell adapters were ligated to the produced double-stranded cDNA. DNA fragments were enriched by PCR using Q5 High-Fidelity 2X Master Mix (New England Biolabs, #M0492S) and sequenced by the Illumina NextSeq 500 system in the Genomics Facility Basel. Sequencing reads were aligned to the human reference genome (GRCh38) using Subread (v1.6.2) using unique mapping (Liao et al., 2013). Expression levels were quantified using the featureCounts function in the Rpackage Rsubread at gene-level (Liao et al.). Normalization across the samples was performed using default parameters in the Rpackage edgeR (Robinson et al., 2010). Differential expression analysis was performed using the exactTest function in the edgeR package. Gene ontology was performed by supplying those differentially expressed genes (adjusted p value < 0.05) to the goana function (Young et al., 2010).

### Single-cell RNA sequencing of human T-cells

Single-cell RNA sequencing was conducted on day 20 of culture for Donor 1 WT (D1 WT) and Donor 1 GSH1 (D1 GSH1) and on day 5 for Donor 2 WT (D2 WT). Single cell 10X libraries were constructed from the isolated single cells following the Chromium Single Cell 3ʹ GEM, Library & Gel Bead Kit v3 (10X Genomics, PN-1000075). Briefly, single cells were co- encapsulated with gel beads (10X Genomics, 2000059) in droplets using Chromium Single Cell B Chip (10X Genomics, 1000074). Final D1 WT, D1 GSH1 and D2 WT libraries were pooled and sequenced on the Illumina NovaSeq platform (26/8/0/93 cycles). Raw sequencing files were supplied to cellranger (v3.1.0) using the count argument under default parameters and the human reference genome (GRCh38-3.0.0). Filtering, normalization and transcriptome analysis was performed using a previously described pipeline in the R package Platypus (Yermanos et al.). Briefly, filtered gene expression matrices from cellranger were supplied as input into the Read10x function in the R package Seurat (Stuart et al., 2019). Cells containing more than 5% mitochondrial genes, or less than 150 unique genes detected were filtered out before using the RunPCA function and subsequent normalization using the function RunHarmony from the Harmony package under default parameters (Korsunsky et al., 2019). Uniform manifold approximation projection was performed with Seurat’s RunUMAP function using the first 20 dimensions and the previously computed Harmony reduction. Clustering was performed by the Seurat functions FindNeighbors and FindClusters using the Harmony reduction and first 20 principal components and the default cluster resolution of 0.5, respectively (Satija et al., 2015). Cluster-specific genes were determined by Seurat’s FindMarkers function for those genes expressed in at least 25% of cells in one of the two groups. Differential genes between samples were calculated using the FindMarkers function from Seurat using the default Wilcoxon Rank Sum Test with Bonferroni multiple hypothesis correction. The source code for the analysis of scRNA-seq data is available at https://github.com/alexyermanos/Platypus.

## Acknowledgements

The authors would like to thank Genomics Facility Basel, particularly Dr Christian Basel, Mrs Katja Eschbach and Mrs Elodie Burcklen and Single Cell Facility of the BSSE department of ETH Zürich, particularly Dr Thomas Horn and Dr Mariangela Di Tacchio. The authors also thank Mr Eric Zigon from the Imaging Core at the Wyss Institute. Additionally, authors thank Mrs Anna Devaux for contributing to experimental work. This study is supported by ETH Research Grants (to STR), Helmut Horten Stiftung (to STR) and Aging and Longevity-Related Research Fund at HMS (to GMC). Erik Aznauryan is the recipient of the Genome Engineer Innovation Grant 2019 from Synthego.

## Author contributions

E.A. and S.T.R designed the study; E.A., A.Y., E.Kap., D.M., G.M.C and S.T.R. contributed to experimental design; E.A. performed experiments; E.Kin. developed the bioinformatic pipeline for GSH identification; E.A. and A.Y analyzed data; G.M.C. and S.T.R. supervised the work; E.A, A.Y. and S.T.R wrote the manuscript with input from authors.

---

**Supplementary table 1. Results of the bioinformatic screen based on established and new criteria**

**Supplementary table 2. DNA sequence of HDR donors used for GSH1 and GSH2 integration**

