## Supplemental figure 1 and 2 for "Discovery and validation of novel human genomic safe harbor sites for gene and cell therapies"

**S1. Gene ontology analysis of HEK293T cells following GSH2 mRuby integration**

**A**

| Term | Ont | N | Up | Down | P.Up | P.Down |
|---|---|---|---|---|---|---|
| response to external biotic stimulus | BP | 443 | 4 | 16 | 0.6143769180 | 0.0002243617 |
| response to other organism | BP | 443 | 4 | 16 | 0.6143769180 | 0.0002243617 |
| ketone body catabolic process | BP | 3 | 2 | 0 | 0.0002689309 | 1.0000000000 |
| negative regulation of viral entry into host cell | BP | 11 | 0 | 3 | 1.0000000000 | 0.0003344033 |
| positive regulation of B cell differentiation | BP | 11 | 0 | 3 | 1.0000000000 | 0.0003344033 |
| response to biotic stimulus | BP | 468 | 4 | 16 | 0.6574891374 | 0.0004134829 |
| positive regulation of gamma-delta T cell activation | BP | 3 | 0 | 2 | 1.0000000000 | 0.0005045283 |
| positive regulation of gamma-delta T cell differentiation | BP | 3 | 0 | 2 | 1.0000000000 | 0.0005045283 |
| positive regulation of synapse structural plasticity | BP | 4 | 2 | 0 | 0.0005345013 | 1.0000000000 |
| granulocyte chemotaxis | BP | 34 | 0 | 4 | 1.0000000000 | 0.0009591469 |
| regulation of gamma-delta T cell activation | BP | 4 | 0 | 2 | 1.0000000000 | 0.0010003837 |
| regulation of gamma-delta T cell differentiation | BP | 4 | 0 | 2 | 1.0000000000 | 0.0010003837 |
| negative regulation of viral life cycle | BP | 59 | 1 | 5 | 0.4326283413 | 0.0010142029 |
| regulation of lamellipodium assembly | BP | 22 | 3 | 0 | 0.0011412228 | 1.0000000000 |
| regulation of synapse structural plasticity | BP | 6 | 2 | 0 | 0.0013196258 | 1.0000000000 |
| epithelial cell proliferation | BP | 227 | 8 | 4 | 0.0014733018 | 0.3448881229 |
| regulation of B cell differentiation | BP | 18 | 0 | 3 | 1.0000000000 | 0.0015462581 |
| regulation of viral entry into host cell | BP | 18 | 0 | 3 | 1.0000000000 | 0.0015462581 |
| granulocyte migration | BP | 39 | 0 | 4 | 1.0000000000 | 0.0016171357 |
| response to wounding | BP | 404 | 11 | 9 | 0.0016423665 | 0.0829742818 |

**B**

| Term | Ont | N | Up | Down | P.Up | P.Down |
|---|---|---|---|---|---|---|
| catalytic activity | MF | 4393 | 60 | 63 | 0.0004425268 | 0.198861176 |
| nucleotide binding | MF | 1938 | 33 | 25 | 0.0004592407 | 0.561260414 |
| nucleoside phosphate binding | MF | 1939 | 33 | 25 | 0.0004636010 | 0.562378980 |
| small molecule binding | MF | 2042 | 34 | 26 | 0.0005547028 | 0.590411411 |
| glucosyltransferase activity | MF | 16 | 0 | 3 | 1.0000000000 | 0.001081704 |
| histone demethylase activity (H3-K4 specific) | MF | 6 | 2 | 0 | 0.0013196258 | 1.000000000 |
| hydro-lyase activity | MF | 40 | 1 | 4 | 0.3188250968 | 0.001778738 |
| purine ribonucleotide binding | MF | 1521 | 26 | 19 | 0.0020434120 | 0.619167125 |
| ribonucleotide binding | MF | 1533 | 26 | 19 | 0.0022831866 | 0.633343282 |
| purine nucleotide binding | MF | 1534 | 26 | 19 | 0.0023042196 | 0.634513905 |
| GTPase activity | MF | 191 | 7 | 0 | 0.0023371550 | 1.000000000 |
| cysteine-type endopeptidase activity involved in apoptotic process | MF | 8 | 2 | 0 | 0.0024326946 | 1.000000000 |
| carbonate dehydratase activity | MF | 6 | 0 | 2 | 1.0000000000 | 0.002458218 |
| purine ribonucleoside triphosphate binding | MF | 1496 | 25 | 19 | 0.0034322371 | 0.588921634 |
| purine ribonucleoside binding | MF | 1502 | 25 | 19 | 0.0036195709 | 0.596262893 |
| purine nucleoside binding | MF | 1504 | 25 | 19 | 0.0036839179 | 0.598698854 |
| ribonucleoside binding | MF | 1505 | 25 | 19 | 0.0037164545 | 0.599914705 |
| nucleoside binding | MF | 1510 | 25 | 19 | 0.0038828266 | 0.605972281 |
| carbohydrate derivative binding | MF | 1711 | 27 | 22 | 0.0051205153 | 0.567117557 |
| structural molecule activity | MF | 493 | 4 | 14 | 0.6973007445 | 0.005219669 |

**C**

| Term | Ont | N | Up | Down | P.Up | P.Down |
|---|---|---|---|---|---|---|
| nucleosome | CC | 75 | 0 | 6 | 1.000000000 | 0.0004338150 |
| protein-DNA complex | CC | 141 | 0 | 8 | 1.000000000 | 0.0005198931 |
| DNA packaging complex | CC | 81 | 0 | 6 | 1.000000000 | 0.0006556767 |
| endoplasmic reticulum lumen | CC | 131 | 2 | 7 | 0.356037244 | 0.0016441876 |
| extracellular region part | CC | 2423 | 21 | 47 | 0.723227120 | 0.0022964254 |
| extracellular region | CC | 2662 | 25 | 50 | 0.571146871 | 0.0031648623 |
| neuron projection | CC | 651 | 14 | 10 | 0.003578165 | 0.3453728522 |
| banded collagen fibril | CC | 8 | 0 | 2 | 1.000000000 | 0.0045104152 |
| fibrillar collagen trimer | CC | 8 | 0 | 2 | 1.000000000 | 0.0045104152 |
| membrane part | CC | 3827 | 50 | 44 | 0.005563888 | 0.8672688832 |
| polymeric cytoskeletal fiber | CC | 445 | 3 | 13 | 0.802157386 | 0.0055882831 |
| supramolecular complex | CC | 445 | 3 | 13 | 0.802157386 | 0.0055882831 |
| supramolecular fiber | CC | 445 | 3 | 13 | 0.802157386 | 0.0055882831 |
| supramolecular polymer | CC | 445 | 3 | 13 | 0.802157386 | 0.0055882831 |
| nuclear nucleosome | CC | 28 | 0 | 3 | 1.000000000 | 0.0056421780 |
| Cul4A-RING E3 ubiquitin ligase complex | CC | 9 | 0 | 2 | 1.000000000 | 0.0057495189 |
| extracellular matrix component | CC | 90 | 0 | 5 | 1.000000000 | 0.0064458127 |
| intrinsic component of membrane | CC | 2956 | 40 | 34 | 0.008817007 | 0.8283156717 |
| matrix side of mitochondrial inner membrane | CC | 1 | 1 | 0 | 0.009537549 | 1.0000000000 |
| integral component of membrane | CC | 2904 | 39 | 34 | 0.011201088 | 0.7947113334 |

### S2. Gene ontology analysis of Jurkat cells following GSH2 mRuby integration

**A**

| Term | Ont | N | Up | Down | P.Up | P.Down |
|---|---|---|---|---|---|---|
| regulation of chondrocyte differentiation | BP | 32 | 0 | 2 | 1 | 0.0008461026 |
| wound healing | BP | 334 | 0 | 4 | 1 | 0.0009106351 |
| cocaine metabolic process | BP | 1 | 0 | 1 | 1 | 0.0013625070 |
| negative regulation of phospholipase A2 activity | BP | 1 | 0 | 1 | 1 | 0.0013625070 |
| neuroblast differentiation | BP | 1 | 0 | 1 | 1 | 0.0013625070 |
| positive regulation of neutrophil apoptotic process | BP | 1 | 0 | 1 | 1 | 0.0013625070 |
| tropane alkaloid metabolic process | BP | 1 | 0 | 1 | 1 | 0.0013625070 |
| regulation of cartilage development | BP | 45 | 0 | 2 | 1 | 0.0016713049 |
| response to wounding | BP | 404 | 0 | 4 | 1 | 0.0018433431 |
| DNA rewinding | BP | 2 | 0 | 1 | 1 | 0.0027232667 |
| fast-twitch skeletal muscle fiber contraction | BP | 2 | 0 | 1 | 1 | 0.0027232667 |
| myoblast migration involved in skeletal muscle regeneration | BP | 2 | 0 | 1 | 1 | 0.0027232667 |
| neutrophil apoptotic process | BP | 2 | 0 | 1 | 1 | 0.0027232667 |
| neutrophil clearance | BP | 2 | 0 | 1 | 1 | 0.0027232667 |
| positive regulation of prostaglandin biosynthetic process | BP | 2 | 0 | 1 | 1 | 0.0027232667 |
| positive regulation of unsaturated fatty acid biosynthetic process | BP | 2 | 0 | 1 | 1 | 0.0027232667 |
| regulation of neutrophil apoptotic process | BP | 2 | 0 | 1 | 1 | 0.0027232667 |
| chondrocyte differentiation | BP | 68 | 0 | 2 | 1 | 0.0037755636 |
| regulation of wound healing | BP | 70 | 0 | 2 | 1 | 0.0039962310 |
| negative regulation of T-helper 2 cell differentiation | BP | 3 | 0 | 1 | 1 | 0.0040822810 |

**B**

| Term | Ont | N | Up | Down | P.Up | P.Down |
|---|---|---|---|---|---|---|
| phospholipase inhibitor activity | MF | 10 | 0 | 2 | 1.000000000 | 7.812861e-05 |
| lipase inhibitor activity | MF | 11 | 0 | 2 | 1.000000000 | 9.541400e-05 |
| calcium-dependent phospholipid binding | MF | 28 | 0 | 2 | 1.000000000 | 6.468814e-04 |
| acetylcholinesterase activity | MF | 1 | 0 | 1 | 1.000000000 | 1.362507e-03 |
| double-stranded DNA-dependent ATPase activity | MF | 1 | 0 | 1 | 1.000000000 | 1.362507e-03 |
| glial cell-derived neurotrophic factor receptor activity | MF | 1 | 0 | 1 | 1.000000000 | 1.362507e-03 |
| enzyme inhibitor activity | MF | 209 | 0 | 3 | 1.000000000 | 2.648948e-03 |
| choline binding | MF | 2 | 0 | 1 | 1.000000000 | 2.723267e-03 |
| cholinesterase activity | MF | 2 | 0 | 1 | 1.000000000 | 2.723267e-03 |
| epidermal growth factor-activated receptor activity | MF | 2 | 0 | 1 | 1.000000000 | 2.723267e-03 |
| polypeptide N-acetylgalactosaminyltransferase activity | MF | 14 | 1 | 0 | 0.003362687 | 1.000000e+00 |
| structural molecule activity | MF | 493 | 0 | 4 | 1.000000000 | 3.801635e-03 |
| phospholipase A2 inhibitor activity | MF | 3 | 0 | 1 | 1.000000000 | 4.082281e-03 |
| cadherin binding involved in cell-cell adhesion | MF | 272 | 0 | 3 | 1.000000000 | 5.555523e-03 |
| protein binding involved in cell-cell adhesion | MF | 279 | 0 | 3 | 1.000000000 | 5.961947e-03 |
| acetylgalactosaminyltransferase activity | MF | 25 | 1 | 0 | 0.005999504 | 1.000000e+00 |
| cadherin binding | MF | 282 | 0 | 3 | 1.000000000 | 6.141495e-03 |
| protein binding involved in cell adhesion | MF | 283 | 0 | 3 | 1.000000000 | 6.202064e-03 |
| endogenous lipid antigen binding | MF | 5 | 0 | 1 | 1.000000000 | 6.795082e-03 |
| exogenous lipid antigen binding | MF | 5 | 0 | 1 | 1.000000000 | 6.795082e-03 |

**C**

| Term | Ont | N | Up | Down | P.Up | P.Down |
|---|---|---|---|---|---|---|
| adherens junction | CC | 629 | 0 | 6 | 1 | 0.0001229812 |
| anchoring junction | CC | 639 | 0 | 6 | 1 | 0.0001341854 |
| sarcolemma | CC | 80 | 0 | 3 | 1 | 0.0001617804 |
| plasma membrane | CC | 2747 | 0 | 10 | 1 | 0.0010935994 |
| focal adhesion | CC | 362 | 0 | 4 | 1 | 0.0012289977 |
| cell-substrate adherens junction | CC | 363 | 0 | 4 | 1 | 0.0012416438 |
| blood microparticle | CC | 39 | 0 | 2 | 1 | 0.0012569701 |
| cell-substrate junction | CC | 365 | 0 | 4 | 1 | 0.0012672117 |
| cell periphery | CC | 2820 | 0 | 10 | 1 | 0.0013583354 |
| cell junction | CC | 1055 | 0 | 6 | 1 | 0.0019737821 |
| endothelial microparticle | CC | 2 | 0 | 1 | 1 | 0.0027232667 |
| neurofilament cytoskeleton | CC | 2 | 0 | 1 | 1 | 0.0027232667 |
| extrinsic component of external side of plasma membrane | CC | 3 | 0 | 1 | 1 | 0.0040822810 |
| intermediate filament cytoskeleton | CC | 80 | 0 | 2 | 1 | 0.0051873829 |
| extracellular space | CC | 566 | 0 | 4 | 1 | 0.0062184195 |
| basal cortex | CC | 5 | 0 | 1 | 1 | 0.0067950824 |
| cornified envelope | CC | 5 | 0 | 1 | 1 | 0.0067950824 |
| extrinsic component of endosome membrane | CC | 5 | 0 | 1 | 1 | 0.0067950824 |
| cell-cell adherens junction | CC | 302 | 0 | 3 | 1 | 0.0074221620 |
| external side of plasma membrane | CC | 106 | 0 | 2 | 1 | 0.0089475814 |
