## Supplemental table 2 for "Discovery and validation of novel human genomic safe harbor sites for gene and cell therapies"

| **Donor name** | **Donor sequence** |
| --- | --- |
| GSH1 CMV-mRuby | CTGCATTTAAGTAGGATTCAATAATTTTAAAGTGCAGGGACAAAATTTCCTCATATGGCTCACTAGCTACATTGCAAATTTCTTGAAATCAGAACACAGAAGTGCAGTCCTGTGCTCGCAATGCAGACTTGCAGGGTGTAGAGGCATAAATGGCTCCAGAGCCAGGGACATGGGTCCAGAGGGGGGTAGTCTCCAGAAGACTCCTTTCGGGCCTATTACCATGCCTCAGAGGTCCAAGTGGGGCATGGTGAATATATTATCCTTTATATTATATTTCTTATATGTCTACAACTGCCACTTGACATTGATTATTGACTAGTTATTAATAGTAATCAATTACGGGGTCATTAGTTCATAGCCCATATATGGAGTTCCGCGTTACATAACTTACGGTAAATGGCCCGCCTGGCTGACCGCCCAACGACCCCCGCCCATTGACGTCAATAATGACGTATGTTCCCATAGTAACGCCAATAGGGACTTTCCATTGACGTCAATGGGTGGACTATTTACGGTAAACTGCCCACTTGGCAGTACATCAAGTGTATCATATGCCAAGTACGCCCCCTATTGACGTCAATGACGGTAAATGGCCCGCCTGGCATTATGCCCAGTACATGACCTTATGGGACTTTCCTACTTGGCAGTACATCTACGTATTAGTCATCGCTATTACCATGGTGATGCGGTTTTGGCAGTACATCAATGGGCGTGGATAGCGGTTTGACTCACGGGGATTTCCAAGTCTCCACCCCATTGACGTCAATGGGAGTTTGTTTTGGCACCAAAATCAACGGGACTTTCCAAAATGTCGTAACAACTCCGCCCCATTGACGCAAATGGGCGGTAGGCGTGTACGGTGGGAGGTCTATATAAGCAGAGCTCTCTGGCTAACTAGAGAACCCACTGCTTACTGGCTTATCGAAATTAATACGACTCACTATAGGGAGACCCAAGCTTGCGGCCGCCACCATGGTGCGGGGTTCTCATCATCATCATCATCATGGTATGGCTAGCATGACTGGTGGACAGCAAATGGGTCGGGATCTGTACGACGATGACGATAAGGATCCGATGGTGTCTAAGGGCGAAGAGCTGATCAAGGAAAATATGCGTATGAAGGTGGTCATGGAAGGTTCGGTCAACGGCCACCAATTCAAATGCACAGGTGAAGGAGAAGGCAATCCGTACATGGGAACTCAAACCATGAGGATCAAAGTCATCGAGGGAGGACCCCTGCCATTTGCCTTTGACATTCTTGCCACGTCGTTCATGTATGGCAGCCGTACTTTTATCAAGTACCCGAAAGGCATTCCTGATTTCTTTAAACAGTCCTTTCCTGAGGGTTTTACTTGGGAAAGAGTTACGAGATACGAAGATGGTGGAGTCGTCACCGTCATGCAGGACACCAGCCTTGAGGATGGCTGTCTCGTTTACCACGTCCAAGTCAGAGGGGTAAACTTTCCCTCCAATGGTCCCGTGATGCAGAAGAAGACCAAGGGTTGGGAGCCTAATACAGAGATGATGTATCCAGCAGATGGTGGTCTGAGGGGATACACTCATATGGCACTGAAAGTTGATGGTGGTGGCCATCTGTCTTGCTCTTTCGTAACAACTTACAGGTCAAAAAAGACCGTCGGGAACATCAAGATGCCCGGTATCCATGCCGTTGATCACCGCCTGGAAAGGTTAGAGGAAAGTGACAATGAAATGTTCGTAGTACAACGCGAACACGCAGTTGCCAAGTTCGCCGGGCTTGGTGGTGGGATGGACGAGCTGTACAAGTAAGAATTCTGCAGATATCCATCACACTGGCGGCCGCTCGAGCATGCATCTAGAGGGCCCTATTCTATAGTGTCACCTAAATGCTAGAGCTCGCTGATCAGCCTCGACTGTGCCTTCTAGTTGCCAGCCATCTGTTGTTTGCCCCTCCCCCGTGCCTTCCTTGACCCTGGAAGGTGCCACTCCCACTGTCCTTTCCTAATAAAATGAGGAAATTGCATCGCATTGTCTGAGTAGGTGTCATTCTATTCTGGGGGGTGGGGTGGGGCAGGACAGCAAGGGGGAGGATTGGGAAGACAATAGCAGGCATGCTGGGGATGCGGTGGGCTCTATGGCATGGCACTAGGACTAAAGGTTGGCCAAAGTACAAGATATTTGTCTTATCTGATGACAACTCTGTGTCCTGGACTCTCTTCCAGAATAAGACCTTTCCTGCAGCACTGCTTGAACTCCTCTTAGCAAGAGGGAAACATGTGAAATGCTACCAAAATAGAATAGAAGTAAATTCTTATTATATTCCTTTGTTCACTCATATCCTGAAGTGCATCAAATCAGGTTTTCTCACCTGTATAATGCTGTATTTTACTTGAGTTGGAATAATTTTGCTTAGAAATAAATAAGTAAAACAGCACCTG |

| **Donor name** | **Donor sequence** |
| --- | --- |
| GSH2 CMV-mRuby | CATTACATCCAAGTTTAGACTCATTGAGCTCTAAATATTTGGGAAAACATATTTAAAGAAATTATATAGGTTTGATCCAAAATCTCTTTGGCACAACTTGAAATATGGGTAATCGTCATGTGAAATTTGTGAATAGGAGAACCCACTGTAGGATACTTAACATAAATCAGCCACATAATTTCTATCACTGATATCCAGGGAATTTCAATGACAAATCTAGTGATAAAAATTGATAAAACATTTTTGATAGTTTTGATACAAGTGAAAGTCATGGGATATCAGACTTAAAAGAAACCTCAGGACATTGATTATTGACTAGTTATTAATAGTAATCAATTACGGGGTCATTAGTTCATAGCCCATATATGGAGTTCCGCGTTACATAACTTACGGTAAATGGCCCGCCTGGCTGACCGCCCAACGACCCCCGCCCATTGACGTCAATAATGACGTATGTTCCCATAGTAACGCCAATAGGGACTTTCCATTGACGTCAATGGGTGGACTATTTACGGTAAACTGCCCACTTGGCAGTACATCAAGTGTATCATATGCCAAGTACGCCCCCTATTGACGTCAATGACGGTAAATGGCCCGCCTGGCATTATGCCCAGTACATGACCTTATGGGACTTTCCTACTTGGCAGTACATCTACGTATTAGTCATCGCTATTACCATGGTGATGCGGTTTTGGCAGTACATCAATGGGCGTGGATAGCGGTTTGACTCACGGGGATTTCCAAGTCTCCACCCCATTGACGTCAATGGGAGTTTGTTTTGGCACCAAAATCAACGGGACTTTCCAAAATGTCGTAACAACTCCGCCCCATTGACGCAAATGGGCGGTAGGCGTGTACGGTGGGAGGTCTATATAAGCAGAGCTCTCTGGCTAACTAGAGAACCCACTGCTTACTGGCTTATCGAAATTAATACGACTCACTATAGGGAGACCCAAGCTTGCGGCCGCCACCATGGTGCGGGGTTCTCATCATCATCATCATCATGGTATGGCTAGCATGACTGGTGGACAGCAAATGGGTCGGGATCTGTACGACGATGACGATAAGGATCCGATGGTGTCTAAGGGCGAAGAGCTGATCAAGGAAAATATGCGTATGAAGGTGGTCATGGAAGGTTCGGTCAACGGCCACCAATTCAAATGCACAGGTGAAGGAGAAGGCAATCCGTACATGGGAACTCAAACCATGAGGATCAAAGTCATCGAGGGAGGACCCCTGCCATTTGCCTTTGACATTCTTGCCACGTCGTTCATGTATGGCAGCCGTACTTTTATCAAGTACCCGAAAGGCATTCCTGATTTCTTTAAACAGTCCTTTCCTGAGGGTTTTACTTGGGAAAGAGTTACGAGATACGAAGATGGTGGAGTCGTCACCGTCATGCAGGACACCAGCCTTGAGGATGGCTGTCTCGTTTACCACGTCCAAGTCAGAGGGGTAAACTTTCCCTCCAATGGTCCCGTGATGCAGAAGAAGACCAAGGGTTGGGAGCCTAATACAGAGATGATGTATCCAGCAGATGGTGGTCTGAGGGGATACACTCATATGGCACTGAAAGTTGATGGTGGTGGCCATCTGTCTTGCTCTTTCGTAACAACTTACAGGTCAAAAAAGACCGTCGGGAACATCAAGATGCCCGGTATCCATGCCGTTGATCACCGCCTGGAAAGGTTAGAGGAAAGTGACAATGAAATGTTCGTAGTACAACGCGAACACGCAGTTGCCAAGTTCGCCGGGCTTGGTGGTGGGATGGACGAGCTGTACAAGTAAGAATTCTGCAGATATCCATCACACTGGCGGCCGCTCGAGCATGCATCTAGAGGGCCCTATTCTATAGTGTCACCTAAATGCTAGAGCTCGCTGATCAGCCTCGACTGTGCCTTCTAGTTGCCAGCCATCTGTTGTTTGCCCCTCCCCCGTGCCTTCCTTGACCCTGGAAGGTGCCACTCCCACTGTCCTTTCCTAATAAAATGAGGAAATTGCATCGCATTGTCTGAGTAGGTGTCATTCTATTCTGGGGGGTGGGGTGGGGCAGGACAGCAAGGGGGAGGATTGGGAAGACAATAGCAGGCATGCTGGGGATGCGGTGGGCTCTATGGTGCTATCAAGTCTGATGTCAGTAATTTTTGGAGGAGACTGAAGTGCAGTGAGACTATCCAAAGTCAGACATGGGGAAAAGCAGAGTCATCCCTCCTAGGCTGCCAAAATCCTCCCCATCCAAGCTCATCCTTGAAGCCCTCACTTAAGACAAAGTTCCTCCCATCCCTTCTGCCTGCTCTGGCATGGTCTGAACCATTTGCCTATTAATTGCCCTGCCTGGTTTCATTTGTTCTTTTTGCTGTATTTAAACTGTGGGAATTCTATTGTTAACCTTTTTCTTGCTCAACTGAACTGTGACA |

| **Donor name** | **Donor sequence** |
| --- | --- |
| GSH1 CMV-LAMB3-T2A-GFP | CTGCATTTAAGTAGGATTCAATAATTTTAAAGTGCAGGGACAAAATTTCCTCATATGGCTCACTAGCTACATTGCAAATTTCTTGAAATCAGAACACAGAAGTGCAGTCCTGTGCTCGCAATGCAGACTTGCAGGGTGTAGAGGCATAAATGGCTCCAGAGCCAGGGACATGGGTCCAGAGGGGGGTAGTCTCCAGAAGACTCCTTTCGGGCCTATTACCATGCCTCAGAGGTCCAAGTGGGGCATGGTGAATATATTATCCTTTATATTATATTTCTTATATGTCTACAACTGCCACTTGACATTGATTATTGACTAGTTATTAATAGTAATCAATTACGGGGTCATTAGTTCATAGCCCATATATGGAGTTCCGCGTTACATAACTTACGGTAAATGGCCCGCCTGGCTGACCGCCCAACGACCCCCGCCCATTGACGTCAATAATGACGTATGTTCCCATAGTAACGCCAATAGGGACTTTCCATTGACGTCAATGGGTGGACTATTTACGGTAAACTGCCCACTTGGCAGTACATCAAGTGTATCATATGCCAAGTACGCCCCCTATTGACGTCAATGACGGTAAATGGCCCGCCTGGCATTATGCCCAGTACATGACCTTATGGGACTTTCCTACTTGGCAGTACATCTACGTATTAGTCATCGCTATTACCATGGTGATGCGGTTTTGGCAGTACATCAATGGGCGTGGATAGCGGTTTGACTCACGGGGATTTCCAAGTCTCCACCCCATTGACGTCAATGGGAGTTTGTTTTGGCACCAAAATCAACGGGACTTTCCAAAATGTCGTAACAACTCCGCCCCATTGACGCAAATGGGCGGTAGGCGTGTACGGTGGGAGGTCTATATAAGCAGAGCTCTCTGGCTAACTAGAGAACCCACTGCTTACTGGCTTATCGAAATTAATACGACTCACTATAGGGAGACCCAAGCTTGCGGCCGCCACCATGAGACCATTCTTCCTCTTGTGTTTTGCCCTGCCTGGCCTCCTGCATGCCCAACAAGCCTGCTCCCGTGGGGCCTGCTATCCACCTGTTGGGGACCTGCTTGTTGGGAGGACCCGGTTTCTCCGAGCTTCATCTACCTGTGGACTGACCAAGCCTGAGACCTACTGCACCCAGTATGGCGAGTGGCAGATGAAATGCTGCAAGTGTGACTCCAGGCAGCCTCACAACTACTACAGTCACCGAGTAGAGAATGTGGCTTCATCCTCCGGCCCCATGCGCTGGTGGCAGTCCCAGAATGATGTGAACCCTGTCTCTCTGCAGCTGGACCTGGACAGGAGATTCCAGCTTCAAGAAGTCATGATGGAGTTCCAGGGGCCCATGCCTGCCGGCATGCTGATTGAGCGCTCCTCAGACTTCGGTAAGACCTGGCGAGTGTACCAGTACCTGGCTGCCGACTGCACCTCCACCTTCCCTCGGGTCCGCCAGGGTCGGCCTCAGAGCTGGCAGGATGTTCGGTGCCAGTCCCTGCCTCAGAGGCCTAATGCACGCCTAAATGGGGGGAAGGTCCAACTTAACCTTATGGATTTAGTGTCTGGGATTCCAGCAACTCAAAGTCAAAAAATTCAAGAGGTGGGGGAGATCACAAACTTGAGAGTCAATTTCACCAGGCTGGCCCCTGTGCCCCAAAGGGGCTACCACCCTCCCAGCGCCTACTATGCTGTGTCCCAGCTCCGTCTGCAGGGGAGCTGCTTCTGTCACGGCCATGCTGATCGCTGCGCACCCAAGCCTGGGGCCTCTGCAGGCCCCTCCACCGCTGTGCAGGTCCACGATGTCTGTGTCTGCCAGCACAACACTGCCGGCCCAAATTGTGAGCGCTGTGCACCCTTCTACAACAACCGGCCCTGGAGACCGGCGGAGGGCCAGGACGCCCATGAATGCCAAAGGTGCGACTGCAATGGGCACTCAGAGACATGTCACTTTGACCCCGCTGTGTTTGCCGCCAGCCAGGGGGCATATGGAGGTGTGTGTGACAATTGCCGGGACCACACCGAAGGCAAGAACTGTGAGCGGTGTCAGCTGCACTATTTCCGGAACCGGCGCCCGGGAGCTTCCATTCAGGAGACCTGCATCTCCTGCGAGTGTGATCCGGATGGGGCAGTGCCAGGGGCTCCCTGTGACCCAGTGACCGGGCAGTGTGTGTGCAAGGAGCATGTGCAGGGAGAGCGCTGTGACCTATGCAAGCCGGGCTTCACTGGACTCACCTACGCCAACCCGCAGGGCTGCCACCGCTGTGACTGCAACATCCTGGGGTCCCGGAGGGACATGCCGTGTGACGAGGAGAGTGGGCGCTGCCTTTGTCTGCCCAACGTGGTGGGTCCCAAATGTGACCAGTGTGCTCCCTACCACTGGAAGCTGGCCAGTGGCCAGGGCTGTGAACCGTGTGCCTGCGACCCGCACAACTCCCTCAGCCCACAGTGCAACCAGTTCACAGGGCAGTGCCCCTGTCGGGAAGGCTTTGGTGGCCTGATGTGCAGCGCTGCAGCCATCCGCCAGTGTCCAGACCGGACCTATGGAGACGTGGCCACAGGATGCCGAGCCTGTGACTGTGATTTCCGGGGAACAGAGGGCCCGGGCTGCGACAAGGCATCAGGCCGCTGCCTCTGCCGCCCTGGCTTGACCGGGCCCCGCTGTGACCAGTGCCAGCGAGGCTACTGCAATCGCTACCCGGTGTGCGTGGCCTGCCACCCTTGCTTCCAGACCTATGATGCGGACCTCCGGGAGCAGGCCCTGCGCTTTGGTAGACTCCGCAATGCCACCGCCAGCCTGTGGTCAGGGCCTGGGCTGGAGGACCGTGGCCTGGCCTCCCGGATCCTAGATGCAAAGAGTAAGATTGAGCAGATCCGAGCAGTTCTCAGCAGCCCCGCAGTCACAGAGCAGGAGGTGGCTCAGGTGGCCAGTGCCATCCTCTCCCTCAGGCGAACTCTCCAGGGCCTGCAGCTGGATCTGCCCCTGGAGGAGGAGACGTTGTCCCTTCCGAGAGACCTGGAGAGTCTTGACAGAAGCTTCAATGGTCTCCTTACTATGTATCAGAGGAAGAGGGAGCAGTTTGAAAAAATAAGCAGTGCTGATCCTTCAGGAGCCTTCCGGATGCTGAGCACAGCCTACGAGCAGTCAGCCCAGGCTGCTCAGCAGGTCTCCGACAGCTCGCGCCTTTTGGACCAGCTCAGGGACAGCCGGAGAGAGGCAGAGAGGCTGGTGCGGCAGGCGGGAGGAGGAGGAGGCACCGGCAGCCCCAAGCTTGTGGCCCTGAGGCTGGAGATGTCTTCGTTGCCTGACCTGACACCCACCTTCAACAAGCTCTGTGGCAACTCCAGGCAGATGGCTTGCACCCCAATATCATGCCCTGGTGAGCTATGTCCCCAAGACAATGGCACAGCCTGTGGCTCCCGCTGCAGGGGTGTCCTTCCCAGGGCCGGTGGGGCCTTCTTGATGGCGGGGCAGGTGGCTGAGCAGCTGCGGGGCTTCAATGCCCAGCTCCAGCGGACCAGGCAGATGATTAGGGCAGCCGAGGAATCTGCCTCACAGATTCAATCCAGTGCCCAGCGCTTGGAGACCCAGGTGAGCGCCAGCCGCTCCCAGATGGAGGAAGATGTCAGACGCACACGGCTCCTAATCCAGCAGGTCCGGGACTTCCTAACAGACCCCGACACTGATGCAGCCACTATCCAGGAGGTCAGCGAGGCCGTGCTGGCCCTGTGGCTGCCCACAGACTCAGCTACTGTTCTGCAGAAGATGAATGAGATCCAGGCCATTGCAGCCAGGCTCCCCAACGTGGACTTGGTGCTGTCCCAGACCAAGCAGGACATTGCGCGTGCCCGCCGGTTGCAGGCTGAGGCTGAGGAAGCCAGGAGCCGAGCCCATGCAGTGGAGGGCCAGGTGGAAGATGTGGTTGGGAACCTGCGGCAGGGGACAGTGGCACTGCAGGAAGCTCAGGACACCATGCAAGGCACCAGCCGCTCCCTTCGGCTTATCCAGGACAGGGTTGCTGAGGTTCAGCAGGTACTGCGGCCAGCAGAAAAGCTGGTGACAAGCATGACCAAGCAGCTGGGTGACTTCTGGACACGGATGGAGGAGCTCCGCCACCAAGCCCGGCAGCAGGGGGCAGAGGCAGTCCAGGCCCAGCAGCTTGCGGAAGGTGCCAGCGAGCAGGCATTGAGTGCCCAAGAGGGATTTGAGAGAATAAAACAAAAGTATGCTGAGTTGAAGGACCGGTTGGGTCAGAGTTCCATGCTGGGTGAGCAGGGTGCCCGGATCCAGAGTGTGAAGACAGAGGCAGAGGAGCTGTTTGGGGAGACCATGGAGATGATGGACAGGATGAAAGACATGGAGTTGGAGCTGCTGCGGGGCAGCCAGGCCATCATGCTGCGCTCGGCGGACCTGACAGGACTGGAGAAGCGTGTGGAGCAGATCCGTGACCACATCAATGGGCGCGTGCTCTACTATGCCACCTGCAAGGAGGGCAGAGGAAGTCTTCTAACATGCGGTGACGTGGAGGAGAATCCCGGCCCTAGCATGGTGAGCAAGGGCGAGGAGCTGTTCACCGGGGTGGTGCCCATCCTGGTCGAGCTGGACGGCGACGTAAACGGCCACAAGTTCAGCGTGTCCGGCGAGGGCGAGGGCGATGCCACCTACGGCAAGCTGACCCTGAAGTTCATCTGCACCACCGGCAAGCTGCCCGTGCCCTGGCCCACCCTCGTGACCACCCTGACCTACGGCGTGCAGTGCTTCAGCCGCTACCCCGACCACATGAAGCAGCACGACTTCTTCAAGTCCGCCATGCCCGAAGGCTACGTCCAGGAGCGCACCATCTTCTTCAAGGACGACGGCAACTACAAGACCCGCGCCGAGGTGAAGTTCGAGGGCGACACCCTGGTGAACCGCATCGAGCTGAAGGGCATCGACTTCAAGGAGGACGGCAACATCCTGGGGCACAAGCTGGAGTACAACTACAACAGCCACAACGTCTATATCATGGCCGACAAGCAGAAGAACGGCATCAAGGTGAACTTCAAGATCCGCCACAACATCGAGGACGGCAGCGTGCAGCTCGCCGACCACTACCAGCAGAACACCCCCATCGGCGACGGCCCCGTGCTGCTGCCCGACAACCACTACCTGAGCACCCAGTCCGCCCTGAGCAAAGACCCCAACGAGAAGCGCGATCACATGGTCCTGCTGGAGTTCGTGACCGCCGCCGGGATCACTCTCGGCATGGACGAGCTGTACAAGTAAAGCGGACTAGTCTAGCAATCAACCTCTGGATTACAAAATTTGTGAAAGATTGACTGGTATTCTTAACTATGTTGCTCCTTTTACGCTATGTGGATACGCTGCTTTAATGCCTTTGTATCATGCTATTGCTTCCCGTATGGCTTTCATTTTCTCCTCCTTGTATAAATCCTGGTTAGTTCTTGCCACGGCGGAACTCATCGCCGCCTGCCTTGCCCGCTGCTGGACAGGGGCTCGGCTGTTGGGCACTGACAATTCCGTGGTGTTTATTTGTGAAATTTGTGATGCTATTGCTTTATTTGTAACCATTCTAGCTTTATTTGTGAAATTTGTGATGCTATTGCTTTATTTGTAACCATTATAAGCTGCAATAAACAAGTTAACAACAACAATTGCATTCATTTTATGTTTCAGGTTCAGGGGGAGATGTGGGAGGTTTTTTAAAGCCATGGCACTAGGACTAAAGGTTGGCCAAAGTACAAGATATTTGTCTTATCTGATGACAACTCTGTGTCCTGGACTCTCTTCCAGAATAAGACCTTTCCTGCAGCACTGCTTGAACTCCTCTTAGCAAGAGGGAAACATGTGAAATGCTACCAAAATAGAATAGAAGTAAATTCTTATTATATTCCTTTGTTCACTCATATCCTGAAGTGCATCAAATCAGGTTTTCTCACCTGTATAATGCTGTATTTTACTTGAGTTGGAATAATTTTGCTTAGAAATAAATAAGTAAAACAGCACCTG |

| **Donor name** | **Donor sequence** |
| --- | --- |
| GSH2 CMV-LAMB3-T2A-GFP | CATTACATCCAAGTTTAGACTCATTGAGCTCTAAATATTTGGGAAAACATATTTAAAGAAATTATATAGGTTTGATCCAAAATCTCTTTGGCACAACTTGAAATATGGGTAATCGTCATGTGAAATTTGTGAATAGGAGAACCCACTGTAGGATACTTAACATAAATCAGCCACATAATTTCTATCACTGATATCCAGGGAATTTCAATGACAAATCTAGTGATAAAAATTGATAAAACATTTTTGATAGTTTTGATACAAGTGAAAGTCATGGGATATCAGACTTAAAAGAAACCTCAGGACATTGATTATTGACTAGTTATTAATAGTAATCAATTACGGGGTCATTAGTTCATAGCCCATATATGGAGTTCCGCGTTACATAACTTACGGTAAATGGCCCGCCTGGCTGACCGCCCAACGACCCCCGCCCATTGACGTCAATAATGACGTATGTTCCCATAGTAACGCCAATAGGGACTTTCCATTGACGTCAATGGGTGGACTATTTACGGTAAACTGCCCACTTGGCAGTACATCAAGTGTATCATATGCCAAGTACGCCCCCTATTGACGTCAATGACGGTAAATGGCCCGCCTGGCATTATGCCCAGTACATGACCTTATGGGACTTTCCTACTTGGCAGTACATCTACGTATTAGTCATCGCTATTACCATGGTGATGCGGTTTTGGCAGTACATCAATGGGCGTGGATAGCGGTTTGACTCACGGGGATTTCCAAGTCTCCACCCCATTGACGTCAATGGGAGTTTGTTTTGGCACCAAAATCAACGGGACTTTCCAAAATGTCGTAACAACTCCGCCCCATTGACGCAAATGGGCGGTAGGCGTGTACGGTGGGAGGTCTATATAAGCAGAGCTCTCTGGCTAACTAGAGAACCCACTGCTTACTGGCTTATCGAAATTAATACGACTCACTATAGGGAGACCCAAGCTTGCGGCCGCCACCATGAGACCATTCTTCCTCTTGTGTTTTGCCCTGCCTGGCCTCCTGCATGCCCAACAAGCCTGCTCCCGTGGGGCCTGCTATCCACCTGTTGGGGACCTGCTTGTTGGGAGGACCCGGTTTCTCCGAGCTTCATCTACCTGTGGACTGACCAAGCCTGAGACCTACTGCACCCAGTATGGCGAGTGGCAGATGAAATGCTGCAAGTGTGACTCCAGGCAGCCTCACAACTACTACAGTCACCGAGTAGAGAATGTGGCTTCATCCTCCGGCCCCATGCGCTGGTGGCAGTCCCAGAATGATGTGAACCCTGTCTCTCTGCAGCTGGACCTGGACAGGAGATTCCAGCTTCAAGAAGTCATGATGGAGTTCCAGGGGCCCATGCCTGCCGGCATGCTGATTGAGCGCTCCTCAGACTTCGGTAAGACCTGGCGAGTGTACCAGTACCTGGCTGCCGACTGCACCTCCACCTTCCCTCGGGTCCGCCAGGGTCGGCCTCAGAGCTGGCAGGATGTTCGGTGCCAGTCCCTGCCTCAGAGGCCTAATGCACGCCTAAATGGGGGGAAGGTCCAACTTAACCTTATGGATTTAGTGTCTGGGATTCCAGCAACTCAAAGTCAAAAAATTCAAGAGGTGGGGGAGATCACAAACTTGAGAGTCAATTTCACCAGGCTGGCCCCTGTGCCCCAAAGGGGCTACCACCCTCCCAGCGCCTACTATGCTGTGTCCCAGCTCCGTCTGCAGGGGAGCTGCTTCTGTCACGGCCATGCTGATCGCTGCGCACCCAAGCCTGGGGCCTCTGCAGGCCCCTCCACCGCTGTGCAGGTCCACGATGTCTGTGTCTGCCAGCACAACACTGCCGGCCCAAATTGTGAGCGCTGTGCACCCTTCTACAACAACCGGCCCTGGAGACCGGCGGAGGGCCAGGACGCCCATGAATGCCAAAGGTGCGACTGCAATGGGCACTCAGAGACATGTCACTTTGACCCCGCTGTGTTTGCCGCCAGCCAGGGGGCATATGGAGGTGTGTGTGACAATTGCCGGGACCACACCGAAGGCAAGAACTGTGAGCGGTGTCAGCTGCACTATTTCCGGAACCGGCGCCCGGGAGCTTCCATTCAGGAGACCTGCATCTCCTGCGAGTGTGATCCGGATGGGGCAGTGCCAGGGGCTCCCTGTGACCCAGTGACCGGGCAGTGTGTGTGCAAGGAGCATGTGCAGGGAGAGCGCTGTGACCTATGCAAGCCGGGCTTCACTGGACTCACCTACGCCAACCCGCAGGGCTGCCACCGCTGTGACTGCAACATCCTGGGGTCCCGGAGGGACATGCCGTGTGACGAGGAGAGTGGGCGCTGCCTTTGTCTGCCCAACGTGGTGGGTCCCAAATGTGACCAGTGTGCTCCCTACCACTGGAAGCTGGCCAGTGGCCAGGGCTGTGAACCGTGTGCCTGCGACCCGCACAACTCCCTCAGCCCACAGTGCAACCAGTTCACAGGGCAGTGCCCCTGTCGGGAAGGCTTTGGTGGCCTGATGTGCAGCGCTGCAGCCATCCGCCAGTGTCCAGACCGGACCTATGGAGACGTGGCCACAGGATGCCGAGCCTGTGACTGTGATTTCCGGGGAACAGAGGGCCCGGGCTGCGACAAGGCATCAGGCCGCTGCCTCTGCCGCCCTGGCTTGACCGGGCCCCGCTGTGACCAGTGCCAGCGAGGCTACTGCAATCGCTACCCGGTGTGCGTGGCCTGCCACCCTTGCTTCCAGACCTATGATGCGGACCTCCGGGAGCAGGCCCTGCGCTTTGGTAGACTCCGCAATGCCACCGCCAGCCTGTGGTCAGGGCCTGGGCTGGAGGACCGTGGCCTGGCCTCCCGGATCCTAGATGCAAAGAGTAAGATTGAGCAGATCCGAGCAGTTCTCAGCAGCCCCGCAGTCACAGAGCAGGAGGTGGCTCAGGTGGCCAGTGCCATCCTCTCCCTCAGGCGAACTCTCCAGGGCCTGCAGCTGGATCTGCCCCTGGAGGAGGAGACGTTGTCCCTTCCGAGAGACCTGGAGAGTCTTGACAGAAGCTTCAATGGTCTCCTTACTATGTATCAGAGGAAGAGGGAGCAGTTTGAAAAAATAAGCAGTGCTGATCCTTCAGGAGCCTTCCGGATGCTGAGCACAGCCTACGAGCAGTCAGCCCAGGCTGCTCAGCAGGTCTCCGACAGCTCGCGCCTTTTGGACCAGCTCAGGGACAGCCGGAGAGAGGCAGAGAGGCTGGTGCGGCAGGCGGGAGGAGGAGGAGGCACCGGCAGCCCCAAGCTTGTGGCCCTGAGGCTGGAGATGTCTTCGTTGCCTGACCTGACACCCACCTTCAACAAGCTCTGTGGCAACTCCAGGCAGATGGCTTGCACCCCAATATCATGCCCTGGTGAGCTATGTCCCCAAGACAATGGCACAGCCTGTGGCTCCCGCTGCAGGGGTGTCCTTCCCAGGGCCGGTGGGGCCTTCTTGATGGCGGGGCAGGTGGCTGAGCAGCTGCGGGGCTTCAATGCCCAGCTCCAGCGGACCAGGCAGATGATTAGGGCAGCCGAGGAATCTGCCTCACAGATTCAATCCAGTGCCCAGCGCTTGGAGACCCAGGTGAGCGCCAGCCGCTCCCAGATGGAGGAAGATGTCAGACGCACACGGCTCCTAATCCAGCAGGTCCGGGACTTCCTAACAGACCCCGACACTGATGCAGCCACTATCCAGGAGGTCAGCGAGGCCGTGCTGGCCCTGTGGCTGCCCACAGACTCAGCTACTGTTCTGCAGAAGATGAATGAGATCCAGGCCATTGCAGCCAGGCTCCCCAACGTGGACTTGGTGCTGTCCCAGACCAAGCAGGACATTGCGCGTGCCCGCCGGTTGCAGGCTGAGGCTGAGGAAGCCAGGAGCCGAGCCCATGCAGTGGAGGGCCAGGTGGAAGATGTGGTTGGGAACCTGCGGCAGGGGACAGTGGCACTGCAGGAAGCTCAGGACACCATGCAAGGCACCAGCCGCTCCCTTCGGCTTATCCAGGACAGGGTTGCTGAGGTTCAGCAGGTACTGCGGCCAGCAGAAAAGCTGGTGACAAGCATGACCAAGCAGCTGGGTGACTTCTGGACACGGATGGAGGAGCTCCGCCACCAAGCCCGGCAGCAGGGGGCAGAGGCAGTCCAGGCCCAGCAGCTTGCGGAAGGTGCCAGCGAGCAGGCATTGAGTGCCCAAGAGGGATTTGAGAGAATAAAACAAAAGTATGCTGAGTTGAAGGACCGGTTGGGTCAGAGTTCCATGCTGGGTGAGCAGGGTGCCCGGATCCAGAGTGTGAAGACAGAGGCAGAGGAGCTGTTTGGGGAGACCATGGAGATGATGGACAGGATGAAAGACATGGAGTTGGAGCTGCTGCGGGGCAGCCAGGCCATCATGCTGCGCTCGGCGGACCTGACAGGACTGGAGAAGCGTGTGGAGCAGATCCGTGACCACATCAATGGGCGCGTGCTCTACTATGCCACCTGCAAGGAGGGCAGAGGAAGTCTTCTAACATGCGGTGACGTGGAGGAGAATCCCGGCCCTAGCATGGTGAGCAAGGGCGAGGAGCTGTTCACCGGGGTGGTGCCCATCCTGGTCGAGCTGGACGGCGACGTAAACGGCCACAAGTTCAGCGTGTCCGGCGAGGGCGAGGGCGATGCCACCTACGGCAAGCTGACCCTGAAGTTCATCTGCACCACCGGCAAGCTGCCCGTGCCCTGGCCCACCCTCGTGACCACCCTGACCTACGGCGTGCAGTGCTTCAGCCGCTACCCCGACCACATGAAGCAGCACGACTTCTTCAAGTCCGCCATGCCCGAAGGCTACGTCCAGGAGCGCACCATCTTCTTCAAGGACGACGGCAACTACAAGACCCGCGCCGAGGTGAAGTTCGAGGGCGACACCCTGGTGAACCGCATCGAGCTGAAGGGCATCGACTTCAAGGAGGACGGCAACATCCTGGGGCACAAGCTGGAGTACAACTACAACAGCCACAACGTCTATATCATGGCCGACAAGCAGAAGAACGGCATCAAGGTGAACTTCAAGATCCGCCACAACATCGAGGACGGCAGCGTGCAGCTCGCCGACCACTACCAGCAGAACACCCCCATCGGCGACGGCCCCGTGCTGCTGCCCGACAACCACTACCTGAGCACCCAGTCCGCCCTGAGCAAAGACCCCAACGAGAAGCGCGATCACATGGTCCTGCTGGAGTTCGTGACCGCCGCCGGGATCACTCTCGGCATGGACGAGCTGTACAAGTAAAGCGGACTAGTCTAGCAATCAACCTCTGGATTACAAAATTTGTGAAAGATTGACTGGTATTCTTAACTATGTTGCTCCTTTTACGCTATGTGGATACGCTGCTTTAATGCCTTTGTATCATGCTATTGCTTCCCGTATGGCTTTCATTTTCTCCTCCTTGTATAAATCCTGGTTAGTTCTTGCCACGGCGGAACTCATCGCCGCCTGCCTTGCCCGCTGCTGGACAGGGGCTCGGCTGTTGGGCACTGACAATTCCGTGGTGTTTATTTGTGAAATTTGTGATGCTATTGCTTTATTTGTAACCATTCTAGCTTTATTTGTGAAATTTGTGATGCTATTGCTTTATTTGTAACCATTATAAGCTGCAATAAACAAGTTAACAACAACAATTGCATTCATTTTATGTTTCAGGTTCAGGGGGAGATGTGGGAGGTTTTTTAAAGCTGCTATCAAGTCTGATGTCAGTAATTTTTGGAGGAGACTGAAGTGCAGTGAGACTATCCAAAGTCAGACATGGGGAAAAGCAGAGTCATCCCTCCTAGGCTGCCAAAATCCTCCCCATCCAAGCTCATCCTTGAAGCCCTCACTTAAGACAAAGTTCCTCCCATCCCTTCTGCCTGCTCTGGCATGGTCTGAACCATTTGCCTATTAATTGCCCTGCCTGGTTTCATTTGTTCTTTTTGCTGTATTTAAACTGTGGGAATTCTATTGTTAACCTTTTTCTTGCTCAACTGAACTGTGACA |
